# Non-canonical ATR signalling via NBS1 phosphorylation propagates fork slowing from stressed to unperturbed nuclear regions

**DOI:** 10.64898/2026.07.09.737522

**Authors:** Jana Krietsch, Ilaria Ceppi, William J. Comstock, Sandra Piquet, Danina Kuster, Francesca Vivalda, Moses Aouami, Larissa Thöny, Stefan Braunshier, Diego Dibitetto, Alessandro A. Sartori, Sophie E. Polo, Marcus B. Smolka, Petr Cejka, Massimo Lopes

## Abstract

DNA replication forks frequently encounter obstacles and remodel into four-way junctions to actively slow DNA fork progression. Fork slowing can also spread to undamaged forks via an ATR-dependent mechanism that remained elusive. Here, using mild genotoxic stress, we show that fork slowing and reversal require full ATR activity, but no canonical ATR activators and signalling partners, defining a non-canonical ATR pathway distinct from origin firing control. Phospho-proteomics in S-phase cells under checkpoint-blind replication stress identified a subset of ATR-dependent phospho-sites, such as S343 on NBS1, the regulatory subunit of the MRN complex. This residue is essential to stimulate MRN exonuclease activity *in vitro* and required in cells for global fork slowing upon mild DNA damage. Ultimately, local UV-C micro-irradiation reveals that ATR-dependent MRN-stimulated resection dampens DNA synthesis at lesions and propagates fork slowing to undamaged chromatin, supporting MRN-mediated ssDNA exposure as a mean to coordinate replication slowdown across the nucleus.

## Introduction

Cells respond to DNA replication stress (RS) by activating signalling pathways and local mechanisms that stabilize replication forks, limit genomic instability, and delay cell cycle progression^1,2^. Central to this response is the kinase ATR, which coordinates multiple pathways that tolerate replication perturbations and suppress genome instability during hyperproliferation and oncogene activation^3,4^. Replication plasticity also enables cancer cells to withstand chemotherapeutic agents that target DNA synthesis or damage the template, making ATR and its associated pathways attractive therapeutic targets^1,5^. Among RS tolerance mechanisms, replication fork reversal - the remodelling of three-way forks into four-way junctions - has emerged as a key genetically controlled response that slows fork progression under stress, thereby promoting fork stability and restart^6, 7^. Surprisingly, fork reversal also occurs at undamaged forks, propagating local replication perturbations into a genome-wide nuclear response^8^. Although this global response requires ATR activity^8^, the underlying molecular mechanisms, ATR targets, and the extent to which key players of the canonical ATR pathway are involved remained unclear.

Accumulation of single-stranded DNA (ssDNA) at stalled replication forks rapidly coated by the ssDNA binding complex RPA initiates ATR signalling^3,9^. Partial replacement of RPA with the recombinase RAD51 promotes efficient fork reversal^10,11^ by priming parental strand reannealing and providing an accessible junction for DNA translocases^12^. Moreover, ssDNA accumulation is directly or indirectly needed for the biochemical activation of these enzymes^13,14,15^. While limited ssDNA is physiologically present at unperturbed forks and increases upon stress due to uncoupling of DNA synthesis and DNA unwinding, stalled forks can also undergo active resection of nascent strands, particularly in cells defective in fork protection^7,16,17^. This latter pathological degradation depends, in most contexts, on prior fork reversal, identifying reversed forks as substrates or recruitment platforms for nucleases. Among the nucleases implicated in nascent strand degradation (e.g., MRE11, DNA2, EXO1)^18–22^, MRE11 - together with RAD50 and NBS1, in the so-called MRN complex - also contributes to physiological RS responses, including checkpoint activation, fork restart, and post-replicative repair^23–26^. However, whether MRN-mediated fork processing may prime fork plasticity under permissive conditions for DNA synthesis was not addressed to date.

Several lines of evidence suggest that the MRN complex is integral to the DNA replication machinery in higher eukaryotes. The complex is essential in cycling but dispensable in quiescent cells^27^, associates with chromatin during S-phase, co-localizes with PCNA, and is enriched on newly synthesized DNA also in unperturbed human cells^28–30^. Taken together, this evidence suggests that the engagement of the MRN complex at replication forks to modulate fork progression, processing and remodelling upon RS may entail rather different regulatory mechanisms from those extensively characterized for its recruitment and activation during double-strand break (DSB) resection^31^.

How ssDNA accumulates at unperturbed replication forks to prime their remodelling into four-way junctions - and thus enable fork reversal in the absence of DNA damage - remains a fundamental unresolved question. Notably, recent work in *Xenopus laevis* suggested that a certain level of nascent strand resection may precede and possibly promote fork remodelling^32^ . In addition to the fact that the MRN complex travels with replication forks, MRE11 activity was shown to mediate active fork slowing upon topoisomerase I poisoning by camptothecin (CPT) or BRCA2 defects, suggesting that – in specific experimental conditions - controlled fork resection may be functionally linked to fork remodelling in promoting active fork slowdown^33,25^.

Here we report that efficient fork reversal requires full ATR activity and yet does not depend on canonical ATR activators (TOPBP1, ETAA1), the effector kinase CHK1, or the downstream target H2AX. Instead, by uncoupling fork reversal from canonical pathway activation, we identify ATR-dependent phosphorylation of NBS1 on S343 as a key event driving active fork slowing during mild replication interference. Notably, S343 is also required for MRN exonuclease activity in DNA end-resection assays and this specific MRN activity is needed for active fork slowing upon mild genotoxic treatments in human cells. Finally, using local UV-C irradiation, we found that ATR-mediated NBS1 S343 phosphorylation promotes local MRN activation at damaged chromatin and propagates fork remodelling and ATR signalling to neighboring, unchallenged chromatin, ultimately throughout the entire nucleus. This ATR-dependent feed-forward mechanism progressively enables nuclear-wide control of fork progression, regardless of a full activation and amplification of the canonical ATR signalling pathway.

## Results

### Full ATR activity is required to promote global fork slowing and reversal, via non-canonical signalling

ATR was shown to limit DNA synthesis and to promote replication fork reversal upon mild replication interference^8^ . However, probing the effect of ATR inactivation on the progression of individual replication forks is technically challenging, as the associated deregulation of origin firing rapidly leads to nucleotide/RPA exhaustion and to a drastic reduction in fork speed, even in the absence of genotoxic treatments^34,35^. To overcome this limitation and assess whether ATR is indeed required for active fork slowing upon mild genotoxic stress, we took advantage of hypomorphic ATR inactivation in HCT116 ATR -/flox cells, which display reportedly a marked reduction in ATR levels but residual activation of canonical ATR signalling^36^ (Fig. 1a and Extended Data Fig. 1a). This residual ATR activity was sufficient to retain normal control of origin firing (Fig. 1b, c) and normal fork progression rates (Extended Data Fig. 1b), as assessed by incorporation of halogenated nucleotides and spread DNA fiber assays^37^. To assess the impact of partial ATR inactivation on active fork slowing and remodelling, we exposed ATR -/flox cells to mild treatment (100 nM) with the topoisomerase I inhibitor camptothecin (CPT), reportedly inducing marked fork slowing and reversal, combined with low levels of replication-associated double-strand breaks (DSBs)^10^; alternatively, we used a mild dose (20 nM) of etoposide (ETP), a topoisomerase II inhibitor that also significantly impacts replication fork progression and remodelling, without inducing detectable DSBs or checkpoint activation^10^ (Fig. 1a, Extended Data Fig. 1c). Strikingly, hypomorphic ATR inactivation completely and reproducibly abolished active fork slowing in response to both treatments (Fig. 1d) and drastically impaired CPT-induced fork reversal (Fig. 1e, f). To directly test the contribution of canonical ATR activators in fork slowing and reversal^3^, we took advantage of HCT116 cells expressing a version of ETAA1 with a mutated ATR-activation domain and thus incapable of activating ATR and carrying an auxin-inducible TOPBP1 degron^38^. In keeping with published results^38^, auxin addition completely impaired canonical ATR signalling upon CPT treatment, as detected by a lack of ATR autophosphorylation (T1989), as well as CHK1/RPA2 phosphorylation (Fig. 1g). Nonetheless, active fork slowing and reversal upon mild CPT or ETP treatments was fully functional in these cells (Fig. 1h, i). Moreover, treatments with two different CHK1 inhibitors - which demonstrably prevent CHK1 activity upon CPT treatment, as assessed by its autophosphorylation^39^ (Fig. 1j) - did not affect the rate of CPT-induced fork reversal, which was instead affected by ATR inhibition, used here as positive control (Fig. 1k). Hence, the role of ATR in replication fork remodelling does not depend on the activity of its canonical signalling partner, i.e. the CHK1 kinase. Differently from ATR defects, efficient inhibition of the ATM kinase did not affect active fork slowing (Extended Data Fig. 1c, d). Moreover, RPE-1 cells carrying the H2AX-S139A mutation^40^ - which prevents efficient phosphorylation of the canonical ATR target H2AX (Extended Data Fig. 1e) - are fully functional for active replication fork slowing upon CPT or ETP treatments (Extended Data Fig. 1f). Overall, these data strongly suggest that full ATR activation is required to mediate active fork slowing and reversal, which is however independent from canonical ATR signalling activators, partners and targets.

**Fig. 1:**
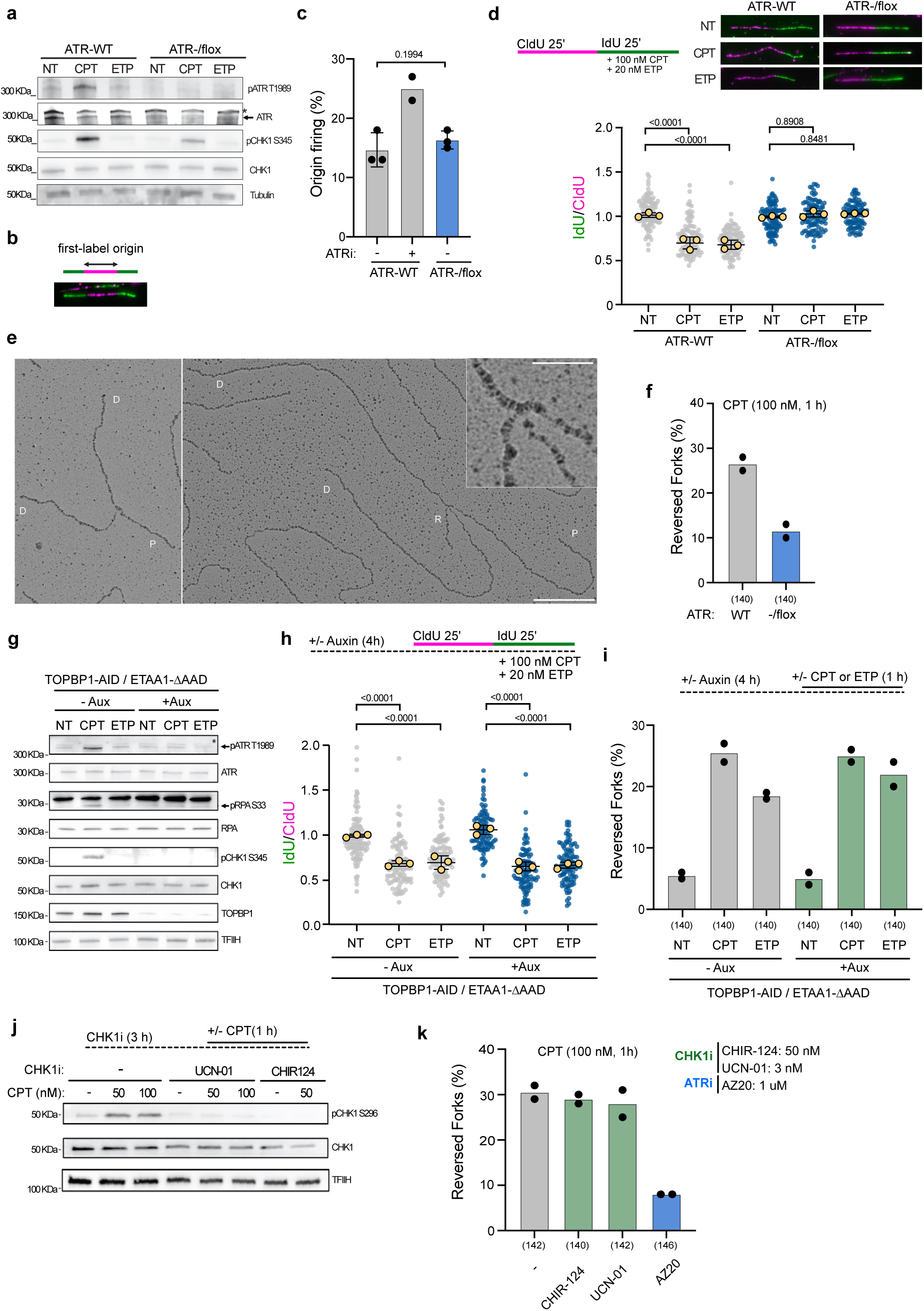
Full ATR activity is required for fork slowing and reversal, but canonical ATR activators, partners and targets are dispensable. a, Validation of HCT116-derived ATR-/flox cell line, hypomorphic for ATR and optionally treated with Campthotecin (CPT, 100 nM, 1 h) or Etoposide (ETP, 20 nM, 1 h) via Western blotting for ATR total protein, autophosphorylated ATR (T1989), phosphorylated CHK1 (S345), CHK1 total protein and Tubulin as loading control. Black arrow indicates specific band, asterisk unspecific band b-d, DNA fiber assay in ATR WT versus hypomorphic conditions from a. b, Representative image of a first label origin. c, Percentage of first-label origins (green-magenta-green molecules) amongst all counted molecules per indicated sample (a minimum of 100 molecules were scored per sample, black dots represent individual biological replicates, grey bars the mean of the replicates). Where indicated, ATR WT cells were pre-treated with ATRi (AZ20, 1 uM, 1 h) as a positive control for elevated levels of newly fired origins. d, top left: Schematic CldU/IdU pulse-labeling protocol used to evaluate fork progression upon 100 nM CPT or 20 nM ETP during the IdU pulse. top right: Representative DNA fiber images. bottom: IdU/CIdU ratio is plotted for a minimum of 100 forks from a single representative experiment (indicated as grey/blue dots). Overlayed yellow dots indicate medians from 3 individual biological replicates. Black line indicates the mean of the medians +/- SD. e, Electron micrographs of representative replication forks from U2OS cells treated with CPT (100 nm, 1 h). left: normal fork; right: reversed fork. parental (P), daughter (D) duplexes, (R) regressed arm; the four-way junction at the reversed fork is in the inset. Scale bar = 200 nm and 40 nm in the inset. f, Frequency (in %) of reversed replication fork molecules isolated from CPT (100 nM, 1 h) treated ATR WT and ATR-/flox cells. Bar graphs depict mean from two independent EM experiments (black dots, respectively). Total number of analyzed molecules in brackets. g, Western blot analysis of HCT116 -derived cell lines containing a truncation in the ATR-activation domain of ETAA1 and proficient (-Aux) or deficient (+Aux) for TOPBP1, treated where indicated with 100 nM CPT or 20 nM ETP for 1 h. Black arrow indicates specific bands. TFIIH serves as loading control. h, DNA fiber assay in ETAA1-mutated (-Aux) and ETAA1 and TOPBP1-deficient (+Aux) HCT116 cells, optionally treated with 100 nM CPT or 20 nM ETP during IdU incorporation. top: Treatment schedule. bottom: IdU/CIdU ratio is plotted for a minimum of 100 forks from a single representative experiment (indicated as grey/ blue dots). Overlayed yellow dots indicate medians from 3 individual biological replicates. Black line indicates the mean of the medians +/- SD. i, Frequency (in %) of reversed replication fork molecules isolated from optionally CPT (100 nM, 1 h) or ETP (20 nM, 1 h) treated HCT116 cells from g and h. Bar graph depicts mean of two independent EM experiments (black dots, respectively). Total number of analyzed molecules in brackets. j, Western blot analysis of U2OS cells treated for 3h where indicated with either UCN-01 (CHK1i, 30 nM) or CHIR124 (CHK1i, 5 nM) and/or CPT (50 or 100 nM, during the last 1 h). k, Frequency (in %) of reversed replication fork molecules isolated from CPT (100 nM, 1 h) treated U2OS cells. Where indicated, cells were pre-treated with a CHK1i or ATRi (positive control). Bar graphs depict mean from two independent EM experiments (black dots, respectively). Total number of analyzed molecules in brackets. c, d, h, p-values were assessed using one-way ANOVA.

### Unbiased screening of non-canonical ATR targets reveals a phospho-site on NBS1 as essential for replication fork slowing

We hence set out to unbiasedly screen for non-canonical ATR targets, seeking for experimental conditions that efficiently induce replication fork slowdown and reversal, with no signs of canonical ATR signalling. Although low dose ETP treatment (20 nM) does not lead to detectable activation of canonical ATR signalling (Fig. 1a, 1g and Extended Data Fig. 1c), fork reversal in these conditions is still clearly dependent on ATR activity (Extended Data Fig. 2a). Hence, we reckoned that this mild ETP treatment may enable identifying ATR-dependent phosphorylation events that are exquisitely required to drive global fork remodelling. To avoid massive DDR signalling associated with chemical synchronization, we turned to FUCCI-U2OS cells^41^ and optimized experimental conditions to sort out unperturbed early S-phase cells (Fig. 2a, b). Sorted cells promptly restarted S-phase progression once re-incubated in conditioned cell culture media, indicating that the isolation procedure as such did not introduce exogenous stress (Extended Data Fig. 2b). We upscaled this sorting and culturing protocol to isolate over fifty million S-phase cells per sample, treated these cells with ETP in presence or absence of ATRi (Fig. 2c) in two independent experiments, and processed these samples for quantitative phospho-proteomic analysis, taking advantage of differential tandem mass tag (TMT) labelling and liquid chromatography coupled to mass-spectrometry (LC–MS). Importantly, while ATR inhibition reportedly leads to fork breakage and ATM signalling upon CPT treatment^42^, this was not the case upon ETP treatment, as neither CHK2-p nor CHK1-p - typically associated with IR-induced DSBs, seen upon ionizing irradiation (IR) - are detectable upon ETP + ATRi treatment (Extended Data Fig. 1c), allowing us to focus the phospho-proteomic screen on basal ATR-dependent events associated with replication fork slowing and remodelling. We focused our attention on S/T-Q sites that were reproducibly decreased upon ATR inhibition in two independent experiments, considering only candidates with Log2(FC) > 0.3 (Supplementary Table 1, Fig. 2d, Extended Data Fig. 2c). As expected, the results of the screen lacked canonical ATR targets such as CHK1. However, they did identify interesting factors involved in nuclear dynamics (e.g. Lamin A/C ^43^) and DNA end resection (e.g. BRCA1^44^ , NBS1^45^) - previously associated with the RS response. Whilst other top hits will be followed up on elsewhere, here we characterized the ATR-dependent phosphorylation of NBS1 - the regulatory subunit of the MRN complex - as in our experimental conditions this protein displayed two independent phosphorylation sites (S343 and S397), which were represented in multiple peptides and reproducibly reduced upon ATR inhibition (Fig. 2d, Extended Data Fig. 2c, Supplementary Table 1). Interestingly, both NBS1 phosphorylation sites identified in our screen mapped within a large intrinsically disordered region (IDR) of the protein. Such regions have been recently reported in multiple DNA damage response factors and their post-translational modification and propensity to form biomolecular condensates were proposed to play key regulatory roles in DDR activation^46^. Phosphorylation of these NBS1 sites by ATM was described in response to DSBs, although their functional contribution to checkpoint signalling and DSB repair is controversial^47,48–51^. Intriguingly, their physiological relevance as non-canonical ATR targets upon mild replication interference - where ATM activity has proven dispensable (Extended Data Fig. 1d) - has not been assessed to date. Profiting from an already commercially available phospho-antibody for the S343 site in NBS1, we validated the ATR-dependency of S343 phosphorylation upon mild ETP treatment, finding that this phosphorylation event is, albeit already present in untreated conditions, comparably elevated upon ETP treatment and fully dependent on ATR activity in both conditions (Fig. 2e).

**Fig. 2:**
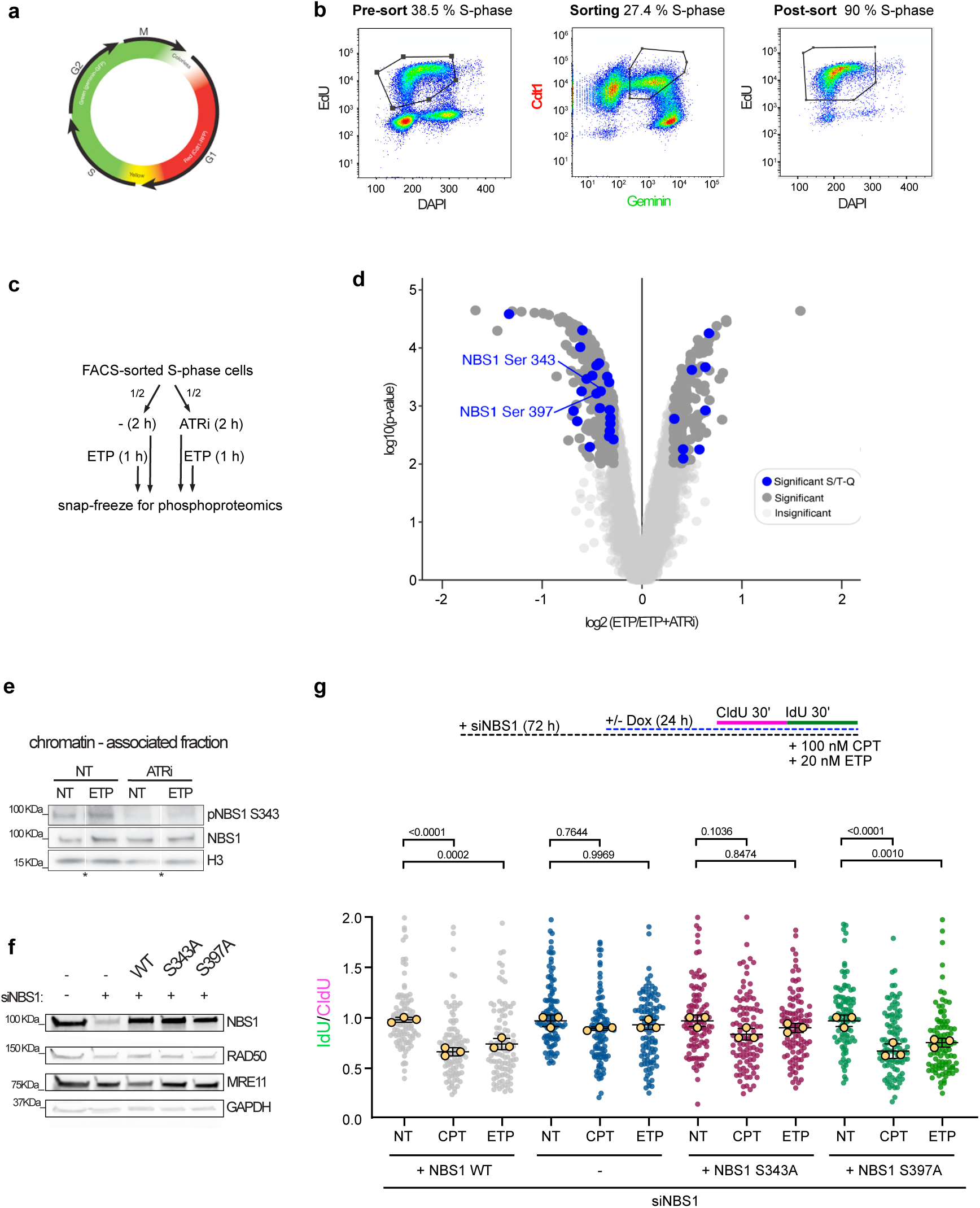
Uncoupling fork slowing from canonical ATR signalling enables phosphoproteomic identification of non-canonical ATR-targets. a, The FUCCI reporter system uses cell cycle–regulated expression of Cdt1-RFP and Geminin-GFP to mark distinct stages, enabling enrichment of S-phase cells without chemical synchronization. b, FACS-based analysis of cells in S-phase in percent. left. EdU incorporation over DAPI, identifying actively replicating cells in the asynchronous cell population prior to sorting. middle: Cdt1/Geminin double-positive population sorted for phospho-proteomics. right: Enrichment of actively replicating cells as identified by EdU incorporation and DAPI staining post sorting. c, Treatment scheme for FACS- sorted S-phase cells: Per 1M sorted cells, the material was divided onto 2 samples. One was left untreated and the other one was preincubated with ATRi (AZ20, 1 uM, 1 h). Subsequently, ETP (20 nM, 1 h) was added to both followed by snap-freezing for phosphoproteomics. d, Volcano plot showing phosphorylation events detected upon ETP treatment and their modulation by ATR inhibition (treatment scheme as in c; significant S/T-Q sites highlighted as dark blue dots). The x-axis indicates the log2 ratio between the two different treatments (ETP and ETP + ATRi) and the y-axis indicates the -log10 probability of each detected phosphorylation event. Two independent biological experiments were performed, yielding comparable results. The screen identified two sites (S343 and S397) on the NBS1 protein. e, Screen target validation using chromatin fractionation followed by Western blotting for S343-specific phosphorylation of NBS1, total chromatin-associated NBS1 and H3 as a loading control. Asterisk indicates where irrelevant samples have been cropped out. f, Western blot analysis of generated U2OS Flip-IN T-Rex cells, enabling the downregulation of endogenous NBS1 protein via siRNA and the parallel doxycycline-induced expression of NBS1 wild type protein (WT) or containing a single point mutation at Serine 343 (S343A) or Serine 397 (S397A). GAPDH serves as loading control. g, DNA fiber assay in U2OS Flip-IN T-Rex cells, downregulating endogenous NBS1 protein with an siRNA and exogenously expressing NBS1-WT or NBS1 S343A or NBS1 S397A, treated where indicated with 100 nM CPT or 20 nM ETP during the IdU pulse. top: Schematic treatment timeline. bottom: IdU/CIdU ratio is plotted for a minimum of 100 forks from a single representative experiment (indicated as grey/ blue/ magenta/ green dots). Overlayed yellow dots indicate medians from 3 individual biological replicates. Black line indicates the mean of the medians +/- SD. g, p-values were assessed using one-way ANOVA.

To assess the functional relevance of NBS1 phosphorylation on S343 and S397 in response to mild genotoxic stress, we derived U2OS stable cell lines where expression of siRNA-resistant, exogenous NBS1 is conditionally induced by doxycycline (DOX) at levels comparable to the endogenous protein (Fig. 2f). Combining DOX addition with siRNA transfection, we can replace endogenous NBS1 with similar levels of wild-type NBS1 or of specific mutant proteins that cannot be phosphorylated on S343 or S397 (S343A, S397A; Fig. 2f). When these cells were tested for the ability to actively slow down fork progression upon CPT or ETP treatment, we observed that both NBS1 depletion (siNBS1) and complementation with the S343A mutant significantly and reproducibly lead to unrestrained fork progression upon treatment (Fig. 2g), while expression of the wild type protein or S397A mutant did restore active fork slowing by CPT/ETP (Fig. 2g). Altogether, these data support the hypothesis that NBS1 and its phosphorylation on S343 are required for efficient replication fork slowdown, while the second phosphorylation event (S397) identified in our phospho-proteomic screen appears irrelevant for these transactions.

### NBS1 S343 regulates reconstituted MRN exonuclease activity, an activity needed in cells for active fork slowdown

Together with MRE11 and RAD50, NBS1 forms the so called MRN complex, heavily studied for its role in DNA end resection in a plethora of biological contexts. Besides its established role in DSB resection and repair^52^, MRE11-dependent activity was shown to contribute to processing and restart of stalled replication forks^23,24,26,53^ and was recently reported to mediate active fork slowing upon mild CPT treatments^25^. Hence, we hypothesized that NBS1 phosphorylation regulates fork progression by modulating MRN-dependent nuclease activity. To test this, we turned to biochemical experiments, taking advantage of substrates and conditions extensively used to study DSB resection^52^.

We purified the MRN complex – containing WT or S343A NBS1 variants – from baculovirus-infected S*podoptera frugiperda* 9 (*Sf*9) insect cells (Extended Data Fig. 3a) to test the functional relevance of the NBS1 S343 residue in regulating MRE11 nuclease activity. We then confirmed that the S343A mutation did not affect MRN complex formation and its binding to DNA, which appeared comparable to that observed for the WT complex in electrophoretic mobility shift assays (Extended Data Fig. 3b, c). In a setup assessing the MRN endonucleolytic activity, phosphorylated CtIP (pCtIP) promotes the clipping of the 5ʹ-terminated DNA strand near protein-blocked DSBs by the MRE11 nuclease within the MRN complex, and this stimulation requires the C-terminal domain of NBS1^52,54^ (Extended Data Fig. 3d). The WT complex – in keeping with published results^54^ - supported the stimulation of MRN endonuclease activity by pCtIP and mutant MRN (S343A) showed comparable activity (Extended Data Fig. 3d, e). Recent work has clarified that – in the context of ssDNA regions arising at stressed replication forks – MRN mostly acts as an exonuclease^55^. Modulation of MRN exonuclease activity by NBS1 has not been studied in detail but appears distinct from the canonical CtIP-dependent activation of MRN endonuclease characterized at DNA ends^55^ . Interestingly, when using a substrate with unprotected ends, MRN-WT exhibited pCtIP-independent exonuclease activity – which was abolished by the single point mutation in MRN (S343A) (Fig. 3a, b), suggesting that the S343 residue in NBS1, or the wider region impacted by the mutation is required for exonucleolytic activity of the MRN complex *in vitro*. Strikingly, inhibiting the exonuclease activity of MRE11 by mirin or the specific MRN exonuclease inhibitor PFM39^56^ completely abolished ETP and CPT-induced fork slowing in U2OS cells (Fig. 3c), hence phenocopying the NBS1 S343A mutation (Fig. 2g). Moreover, the frequency of fork reversal was markedly reduced in response to CPT when MRE11 exonuclease activity was inhibited by either inhibitor (Fig. 3d). Altogether these results suggest that non-canonical ATR-mediated NBS1 phosphorylation at S343 stimulates MRE11 exonuclease activity to actively induce replication fork reversal and slowdown upon mild, systemic treatments.

**Fig. 3:**
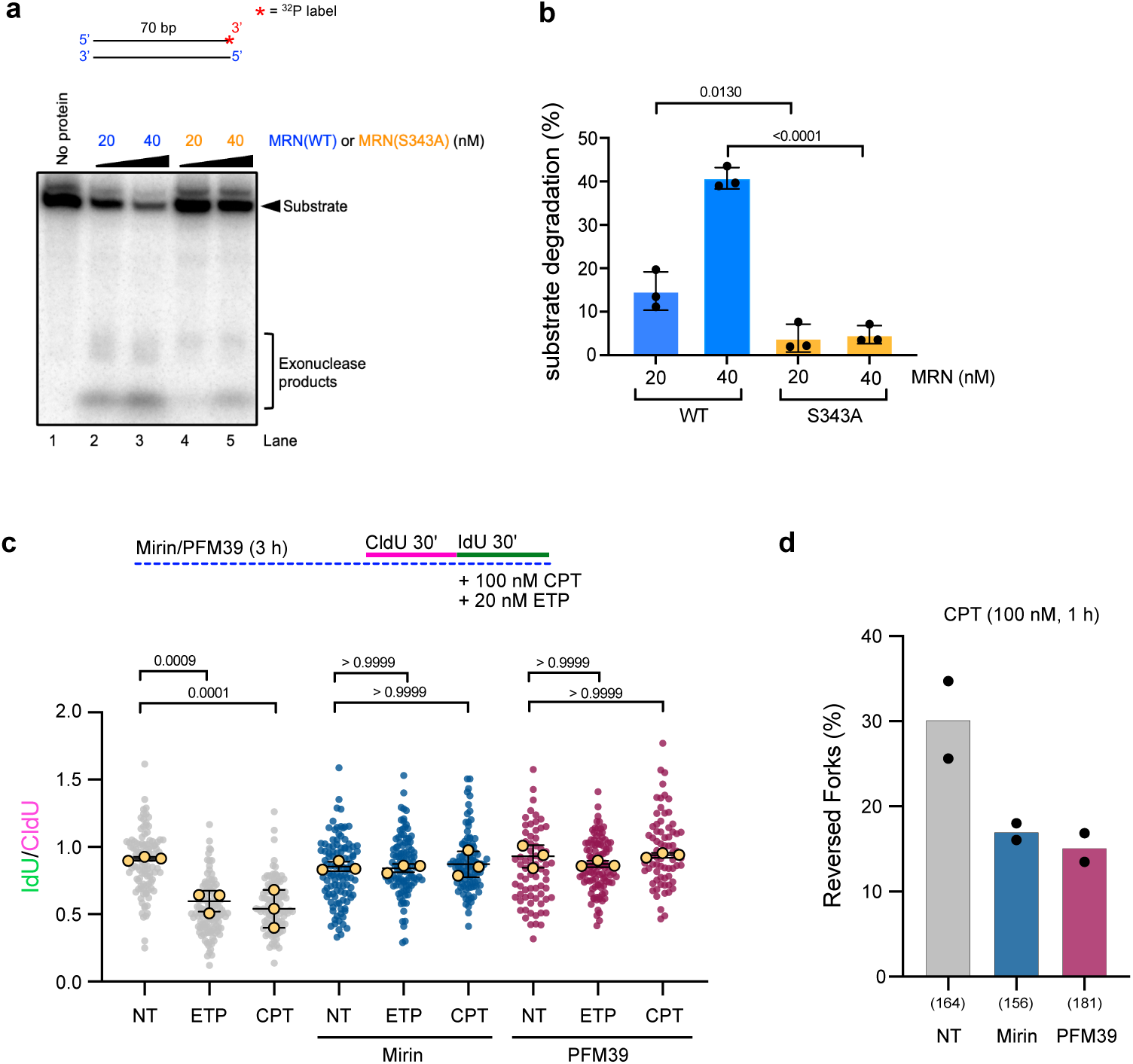
A single amino acid substitution in NBS1 compromises MRN exonuclease activity *in vitro* and inhibiting this activity impairs efficient fork slowing and reversal *in vivo.* a, Biochemical assessment of substrate degradation by recombinant MRN complex containing either the wild type NBS1 protein (MRN WT) or NBS1 mutated on Serine 343 (MRN S343A). top: Schematic depiction of the ^32^P-labeled dsDNA with unprotected ends, used to asssess exonucleolytic activity of purified MRN. bottom: Representative radiograph visualizing substrate degradation by increasing amounts of the indicated protein complexes. b, Substrate degradation in percent in the indicated conditions. Bar graphs display mean +/- SEM, black dots represent means of three independent biological replicates. c, DNA fiber assay in U2OS cells, optionally pre-treated with two resection inhibitors (Mirin, MRE11i, 50 µM, 2 h pre-treatment, 3 h total; PFM39, MRE11exoi, 50 uM, 2 h pretreatment, 3 h total) and treated where indicated with 100 nM CPT or 20 nM ETP during the second label. top: Schematic treatment timeline. bottom: IdU/CIdU ratio is plotted for a minimum of 100 forks from a single representative experiment (indicated as grey/ blue/ magenta dots). Overlayed yellow dots indicate medians from 3 individual biological replicates. Black line indicates the mean of the medians +/-SD. b, c p-values were assessed using one-way ANOVA. d, Frequency (in %) of reversed replication forks isolated from CPT (100 nM, 1 h) treated U2OS cells, optionally treated with resection inhibitors (Mirin, MRE11i, 50 µM, 2 h pre-treatment, 3 h total; PFM39, MRE11exoi, 50 uM, 2 h pretreatment, 3 h total). Grey and blue bar graphs depict mean from two independent EM experiments (black dots, respectively). Total number of analyzed molecules in brackets.

### Localized UV-C DNA lesions reduce replicative DNA synthesis at lesion sites and throughout undamaged subnuclear areas

Active fork slowing and remodelling is a global, ATR-dependent response, which can extend to unchallenged replication forks upon mild and systemic genotoxic treatments^8^. However, to which extent this RS response can physically spread from local DNA lesions to undamaged chromatin and possibly throughout intact nuclei has not been directly assessed. Hence, we set out to investigate these signalling mechanisms and to assess a possible dependency on non-canonical ATR-signalling and MRN activity.

To limit DNA damage to defined subnuclear locations in live nuclei, we used a short (3s) UV-C pulse to irradiate cells covered by a micro-pore filter, knowingly shielding 98 % of UV-C light^57^ (Fig. 4a and Extended Data Fig. 4a), allowing for efficient introduction of photolesions (cyclobutene pyrimidine dimers, CPD) in subnuclear areas (Fig. 4b). Cells were left to recover for 30 min and DNA synthesis was measured by adding EdU in the final minutes of this recovery phase. Using a FIJI-based image analysis home-made pipeline, subnuclear masks were generated to quantify mean fluorescence intensities. Measurements were obtained from the entire nuclei of undamaged cells (lacking a CPD signal), and from irradiated cells either specifically within CPD-positive (damaged) regions, or within undamaged regions, defined as the whole nucleus excluding the CPD-positive area (Extended Data Fig. 4b). Filter-shielded areas remained free of UV-C induced CPD lesions, comparable to a non-irradiated control (Fig. 4b, c). Investigating the rate of replicative DNA synthesis with nucleotide analog (EdU) incorporation in this setup expectedly showed a significant slowdown within the damaged areas when compared to non-irradiated cells. Interestingly the undamaged regions of both U2OS and RPE-1 cells overall exhibited a significant reduction in DNA synthesis (Fig. 4d, e), indicating that DNA synthesis is actively reduced throughout the nucleus. To validate the presence of a globally integrated RS response, we made use of another experimental setup, where laser micro-irradiation with a UV-C photomanipulation unit was directly coupled to confocal microscope (Fig. 4f, Extended Data Fig. 4c). Also here, the rate of DNA synthesis was markedly reduced within the damaged region of laser-irradiated U2OS cells. Interestingly, this optical sectioning imaging modality frequently revealed reduction of EdU signal beyond the damaged regions at the time point selected for analysis (Fig. 4g, white arrow) and confirmed an overall significant reduction of EdU mean intensity in undamaged nuclear regions when compared to undamaged nuclei (Fig. 4g, h, i).

**Fig. 4:**
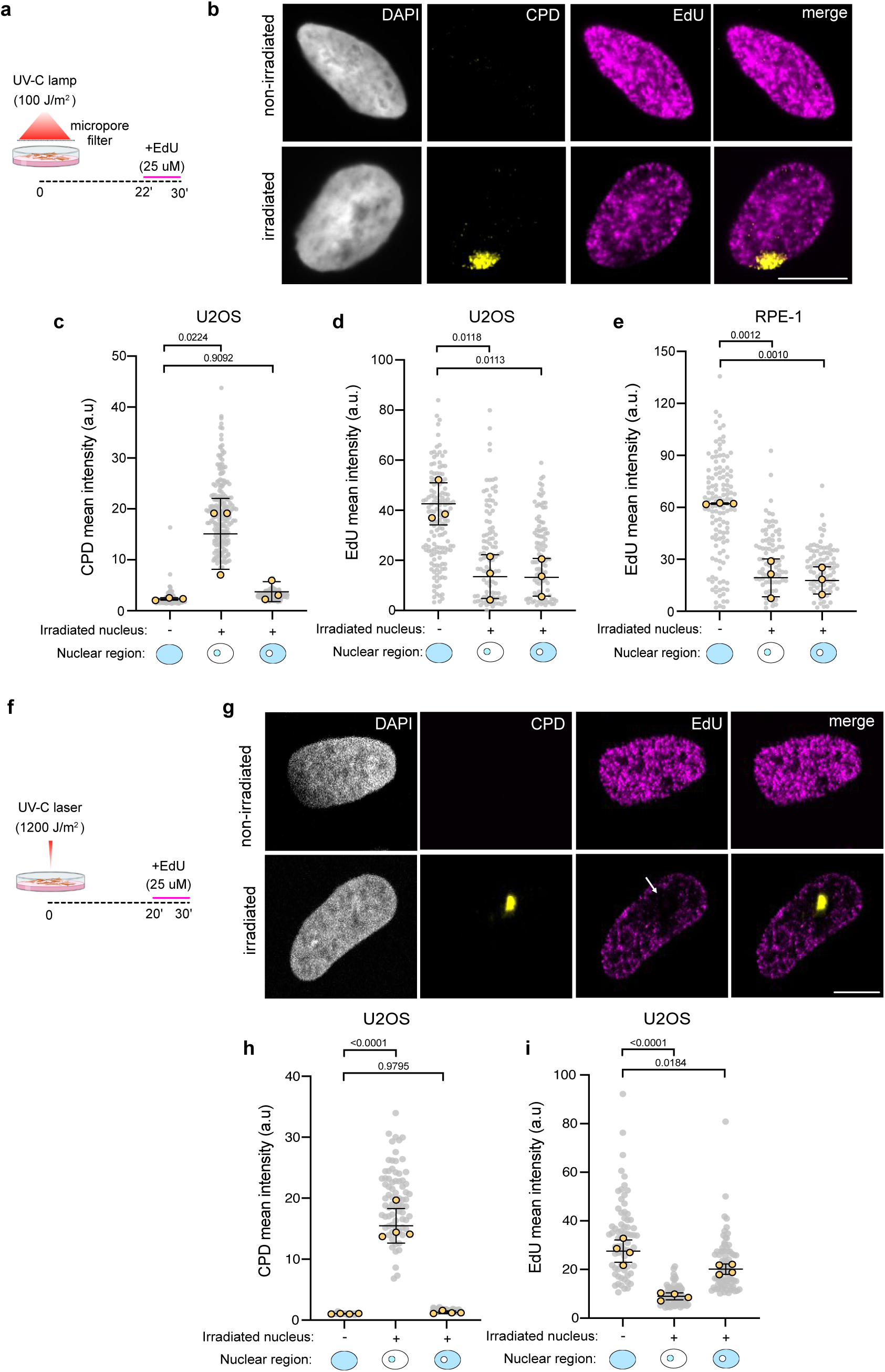
Local DNA lesions reduce DNA synthesis at lesion sites and throughout undamaged subnuclear areas. a, Schematic experimental setup: 100J/m^2^ UV-C irradiation was applied onto a monolayer of cells through a micropore filter. Irradiated or non-irradiated control cells were allowed to recover for 30 mins, with a short (8mins) EdU pulse at the end (see Extended Data Fig. 4a and methods section for more details on the setup and image analysis pipeline). b, Representative widefield microscopy images from an unirradiated U2OS nucleus and one that received UV-C irradiation through the filter pore. Total DNA was counterstained with DAPI (grey), UV-C-induced DNA damage with an antibody against CPD photoproducts (yellow), EdU was clicked with a fluorophore (magenta). Scale bar = 10 µm. c, Using the setup from A in U2OS cells, CPD mean intensity was measured in the corresponding nuclear regions (total nucleus, damaged area alone, undamaged area alone, repectively, indicated as light blue below the graph). d, EdU mean intensity measurments in the corresponding regions assessed for CPD intensity in C. e, EdU mean intensity measurements in the depicted subnuclear regions in RPE-1 cells. c-e, Individual nuclei from one representative experiment are depicted in grey, medians from 3 biological replicates in yellow. Black line marks mean of the medians +/- std dev. P-values were assessed with one-way ANOVA. f, Schematic experimental setup: Targeted UV-C micro-irradiation with a 266nm photomanipulation unit coupled to a confocal microscope. Target cells were irradiated and left to recover for a total time of 30 mins. A short EdU pulse was provided during the last 10 mins of recovery (see Extended Data Fig. 4b and methods section for details on the setup and image analysis). g, Representative confocal microscopy images from an unirradiated U2OS nucleus and one that received UV-C irradiation by the laser. Total DNA was counterstained with DAPI (grey), UV-C-induced DNA damage with an antibody against CPD photoproducts (yellow), EdU was clicked with a fluorophore (magenta). White arrow points to EdU drop surrounding the lesion site. Scale bar = 10 µm. h, Using the setup from f in U2OS cells, CPD mean intensity was measured in the corresponding nuclear regions (total nucleus, damaged area alone, undamaged area alone, repectively, indicated as light blue below graphs). i, EdU mean intensity measurments in the depicted subnuclear regions in U2OS cells from h. h-i, Individual nuclei from one representative experiment are depicted in grey, medians from 3 biological replicates in yellow. Black line marks mean of the medians +/- std dev. P-values were assessed with one-way ANOVA.

Together, both setups establish reduced DNA synthesis as an active process that is globally propagated beyond directly damaged chromatin, providing an experimental pipeline to test the genetic dependencies of this signalling mechanism.

### Global RS response propagation is mediated by ATR and MRE11 exonuclease activity

ATR activity is essential for active fork slowdown and remodelling (Fig. 1d, f, Extended Data Fig. 2a); however, whether this entails direct modulation of replication fork progression in undamaged areas of the nucleus remains an important unanswered question. Reverting to ATR -/flox HCT116 cells, we found that hypomorphic ATR activity (Fig. 1a-f, Extended Data Fig. 1a) – which retains competence for origin firing regulation (Fig. 1c) – is insufficient to mediate the observed slowdown of DNA synthesis in both UV-damaged and undamaged areas of the nucleus (Fig. 5a), clearly linking the ATR response to active slowdown of ongoing replication forks throughout the nucleus. Expectedly, full inhibition of ATR enzymatic activity also abolished the propagation of replication fork slowdown within and beyond the UV-damaged regions (Fig. 5b). To address whether ATR-dependent phosphorylation of NBS1 on S343 was implicated in the propagation of the RS response beyond locally irradiated regions, we validated and used in immunofluorescence after UV-C micro-irradiation the pNBS1 antibody originally developed for Western blotting (Fig. 2e, Extended Data Fig. 5a). Strikingly, we found elevated pNBS1 levels on both damaged and undamaged chromatin, when compared to undamaged cells (Fig. 5c, d). As the S343 site is essential for the exonucleolytic activity of reconstituted MRN (Fig. 3a, b) and this activity is needed for global replication fork slow down upon mild systemic treatments (Fig. 3c), we tested directly if MRE11 exonuclease activity was also needed for the reduction in DNA synthesis observed in regions not directly experiencing UV-C damage. Indeed, pre-treatment of cells with the MRE11 exonuclease inhibitor PFM39 prevented the global reduction of DNA synthesis upon localized UV-C irradiation (Fig. 5e), strongly suggesting that fork resection is a pre-requisite for active fork slowdown in unperturbed regions. Single-molecule, population-based studies have repeatedly linked active replication fork slowing upon mild replication interference to replication fork reversal^7^ a transaction requiring the recruitment of the RAD51 recombinase^10,12^ to forks and active engagement of specialized translocases such as SMARCAL1, which is a stable component of the replisome^58,13^. Hence, to directly assess the functional relevance of replication fork reversal for the global fork slowdown in the undamaged nuclear compartments, we exploited our setup in Fig. 4a and combined it with efficient, acute and transient downregulation of SMARCAL1. For this purpose, we generated an HCT116 cell line where SMARCAL1 was endogenously tagged with Halo at the N-terminal end (Halo-SMARCAL1 cells). Treating these cells with Halo-PROTAC3 (HP3) for 24h yielded over 95% reduction of the protein levels outperforming siRNA-mediated downregulation in this setup (Extended Fig. 5b). To validate the cell line, we performed DNA fiber experiments. Herein, Halo-SMARCAL1 cells showed fork slowdown in response to mild Hydroxyurea (HU) treatment (50 µM) compatible with fork progression; SMARCAL1 depletion by HP3 in those cells led to unrestrained fork progression validating defective fork reversal (Extended Fig. 5c). Strikingly, using this cell line in the micro-irradiation setup (Fig. 4a), we found that both the local and global reduction of DNA synthesis were dependent on SMARCAL1 protein levels, indicating that replication fork reversal is induced not only at DNA damage sites but also in undamaged nuclear regions (Fig. 5f).

**Fig. 5:**
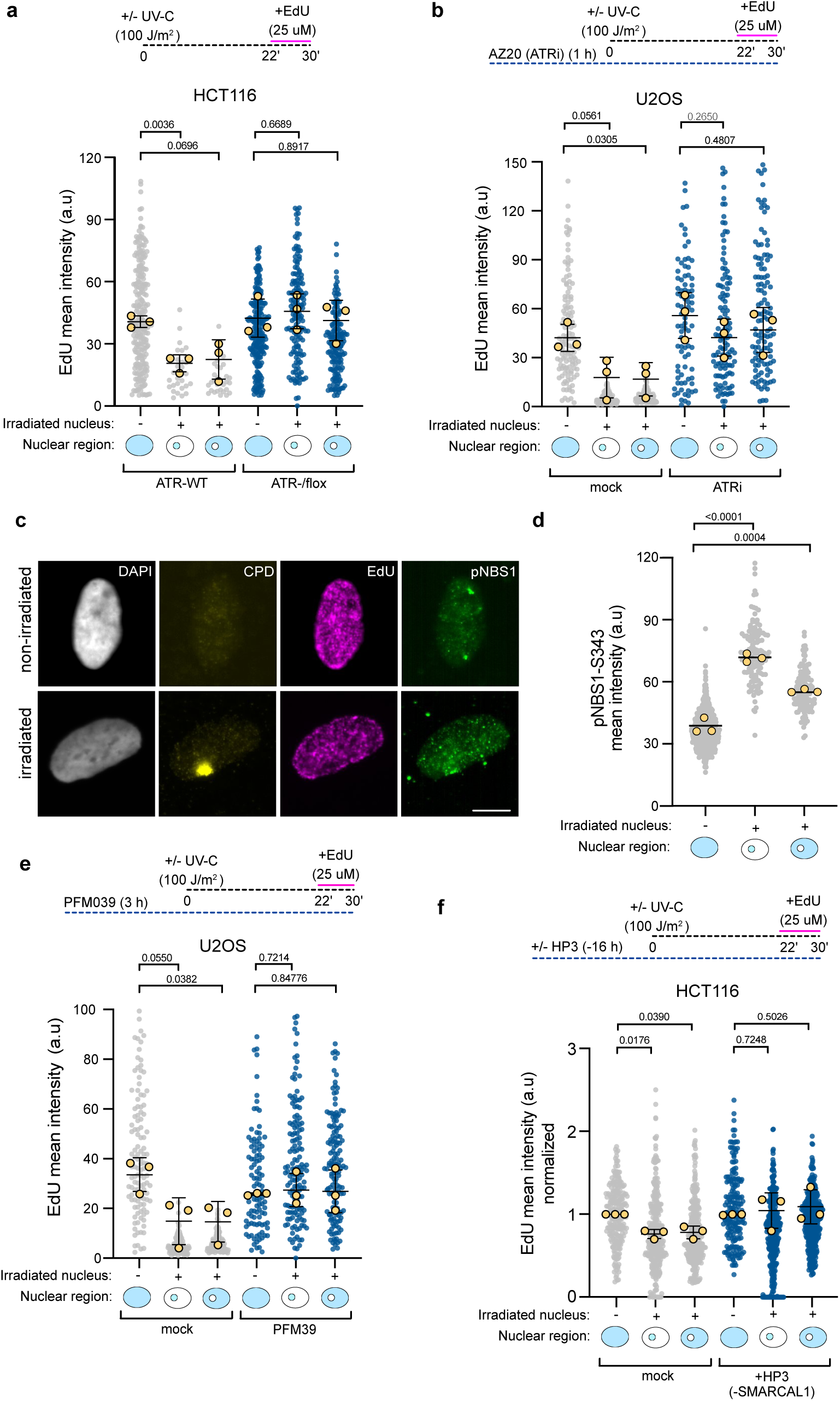
Global replication slowdown after local damage relies on ATR, MRN exonuclease activity and fork reversal. a, EdU mean intensity measurements (experimental setup as in Fig. 4a and Extended Data Fig. 4a,b) in hypomorphic ATR conditions (HCT116-derived ATR-/flox cells, Fig. 1a-f, Extended Data Fig. 1a,b). top: Schematic timeline of the experiment. bottom: EdU mean intensity was measured in the indicated nuclear regions (total nucleus, damaged area alone, undamaged area alone, repectively, indicated as light blue below the graph). b, U2OS cells were optionally treated with ATRi (AZ20, 1 uM, 30 mins pre-treatment and 30 mins during recovery) for EdU mean intensity measurments as in a. c, Representative widefield microscopy images of pNBS1-S343 detected in damaged and undamaged regions (setup as in a, green). d, pNBS1 mean intensity was measured in the indicated nuclear regions (total nucleus, damaged area alone, undamaged area alone, repectively, indicated as light blue below the graph). e, U2OS cells were optionally treated with an MRN exonuclease inhibitor (PFM39, 50 uM, 2 h pre-treatment and maintained during recovery from UV-C) for EdU mean intensity measurements as in A. f, HCT116 Smarcal1 Halo Protac cells (Supplementary Figure 5 a-c) were used in a setup as in A to optionally deplete the translocase SMARCAL1 16h prior to UV-C irradiation. a,b,d-f, Individual nuclei from one representative experiment are displayed as grey/ blue dots, medians from 3 biological replicates in yellow. Black line marks mean of the medians +/- std dev. P values were calculated using Welch’s-corrected two-tailed t test.

Taken together, our results suggest that local UV-C irradiation triggers non-canonical ATR phosphorylation of pNBS1 in damaged and undamaged chromatin, inducing MRE11-dependent exonuclease activity, which is required for active fork slowdown and reversal throughout the nucleus.

## Discussion

Our data shed light on a non-canonical ATR function, modulating replication fork progression and remodelling beyond damaged nuclear areas. Extending previous evidence obtained with ATR inhibitors^8^, we demonstrate that hypomorphic ATR inactivation - permissive for origin firing control - suffices to abolish active fork slowing and efficient fork reversal in human cells. ATR-mediated fork slowing operates independently of TOPBP1, ETAA1, CHK1 and H2AX, providing direct genetic support to the previous finding that RS can induce fork slowing and reversal without detectable activation of canonical ATR signalling markers, such as CHK1- or RPA phosphorylation^10,59^. Our data do not exclude that canonical ATR targets may play a role fine-tuning fork slowing and remodelling in conditions that induce full ATR activation. Moreover, other PI3 kinases (ATM, DNA-PK) were shown to contribute to reversed fork formation and stability at endogenous replication obstacles or upon acute or chronic genotoxic conditions^60,61^. Upon DSB-inducing treatments, NBS1-S343 phosphorylation is mostly ATM-dependent^51,62,47^, but may still contribute to modulate fork progression and remodelling, in parallel to classical DSB signalling. Overall, our data suggest that nuclear propagation of fork slowdown and reversal can also occur in checkpoint-blind conditions, but full activation of canonical checkpoints likely contributes to modulate fork stalling, protection and restart upon particularly harsh genotoxic stress.

Additional ATR/Mec1 functions – even extending beyond the RS response – have been described in yeast and human cells as independent of CHK1 and/or its canonical activators^63–66,67^ supporting the concept that key ATR activities can rely on alternative activation and signalling mechanisms. As suggested by studies in yeast, ssDNA/RPA binding by ATR/ATRIP (Mec1/Ddc2) triggers sufficient kinase activation to execute key replication-related functions, which are genetically distinct from additional signalling roles of the kinase^67,68^. In this perspective, canonical ATR activators and markers constitute a signal amplification layer that is commonly triggered by genotoxic stress but is not essential for all ATR functions in the RS response. Upon mild genotoxic treatments compatible with fork progression, we envision that local ATR recruitment and activation at the subset of replication forks directly challenged by DNA lesions is sufficient to induce NBS1 phosphorylation in proximal chromosomal regions, thereby engaging MRN exonuclease activity at neighboring replication forks that are not directly damaged (Fig. 6). In a feed-forward mechanism, controlled generation of new ssDNA at replication forks could then propagate ATR activation to undamaged DNA, progressively extending ATR signalling to additional factories and ultimately throughout the entire nucleus (Fig. 6). Consistent with this view, it has been shown in a different context that the MRN complex can act both upstream, and downstream of ATR^69^. Depending on the type and level of DNA damage experienced by the cells and on the molecular features of the DNA repair intermediates, this signalling cascade may or may not be associated with canonical markers of ATR activation and signal amplification, which are however not required for global fork slowdown and remodelling.

**Fig. 6:**
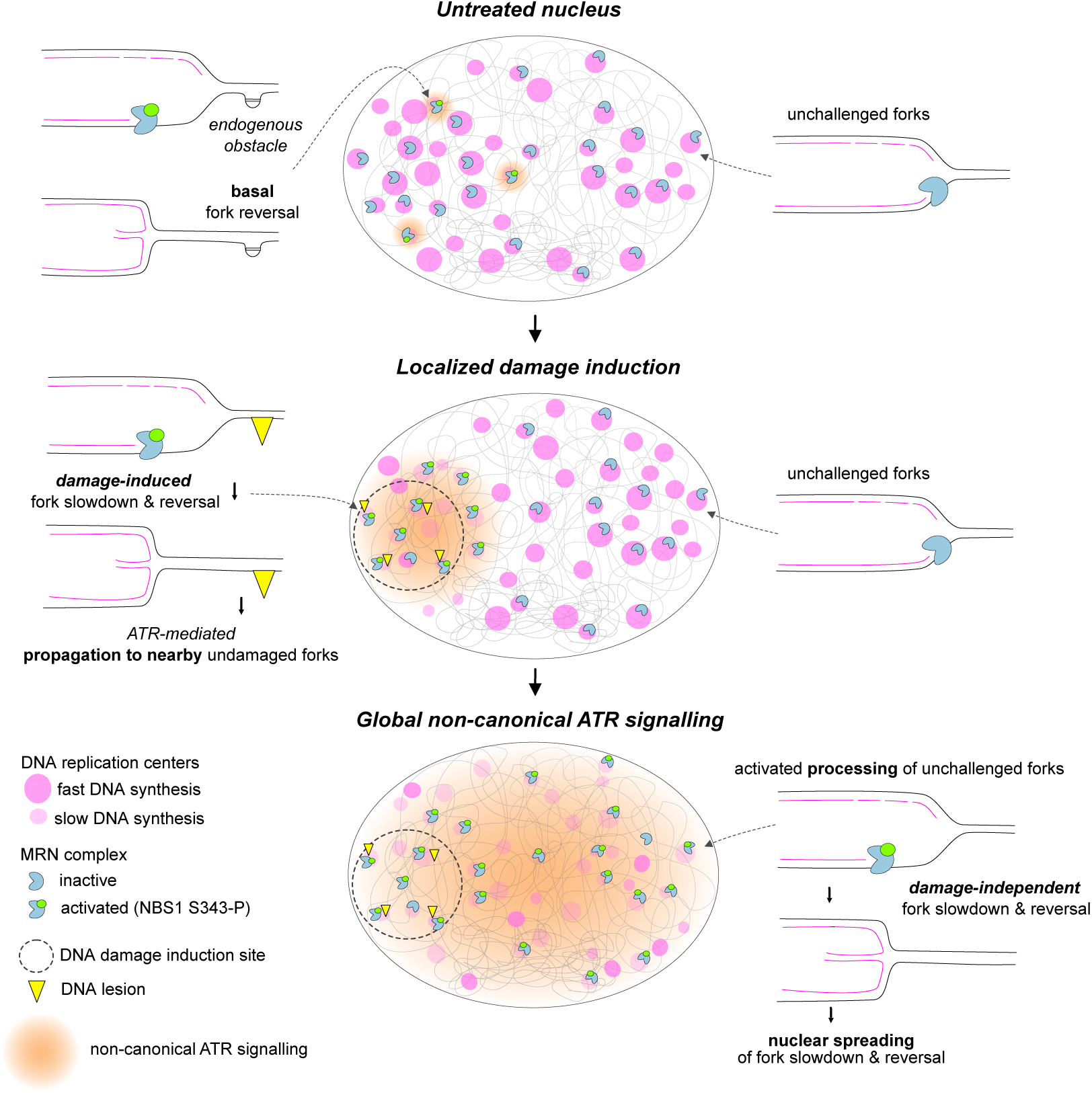
Working model for nuclear propagation of active fork slowing via non-canonical ATR signalling. In untreated nuclei, endogenous obstacles locally induce ATR-dependent NBS1 S343 phosphorylation (green dot), leading to activation (blue pac-man containing green dot) of the otherwise inactive MRN complex (blue pac-man alone) associated with DNA replication centers (magenta circles). Upon localized damage induction, ATR-mediated NBS1 S343 phosphorylation promotes MRN exonuclease activity directly at DNA lesions (yellow triangles), leading to reduced DNA synthesis and reversal at the corresponding replication centers (light magenta, smaller circles). ATR signals this event to directly neighboring, unchallenged chromatin, leading to NBS1 S343 phosphorylation, MRN activation, fork slowing and remodelling at nearby undamaged forks – which in turn leads to more ATR signalling. Ultimately, this stepwise, feed-forward mechanism leads to global non-canonical ATR signalling promoting nuclear spreading of fork slowdown & reversal throughout undamaged chromatin.

It may seem counterintuitive that potentially dangerous transactions are actively induced also at forks that are not directly experiencing any issue, especially if DNA synthesis is anyway limited globally by preventing origin firing. We note, however, that protection and restart of remodelled forks are efficient and finely tuned mechanisms, while pathological processing of reversed forks is only observed in certain genetic contexts. Strikingly, fork reversal and restart are widespread even in key cells giving rise to entire organisms (i.e. ESCs), in the absence of exogenous DNA damage^70^. Moreover, although origin control would *per se* prevent potentially dangerous DNA synthesis in undamaged regions, it does require full activation of canonical ATR signalling, which may not be intrinsic to all genotoxic treatments, and it would not protect forks that are already active at the time of local damage induction. Interestingly, in the context of transcription - where issues with ongoing events may be arguably better tolerated than during genome duplication - global control of elongation and restart is already well established, besides inhibition of new initiation events, to globally fine-tune transcription in trans to the lesions, even in response to minimal, localized DNA damage^71,72,73,74^.

Multiple key drivers of fork reversal – i.e. RAD51 chromatin loading and strand exchange activity^10,12,13^, SMARCAL1 recruitment via RPA binding^14^ , ZRANB3 recruitment via PCNA polyubiquitination^15,75^, activation of the HLTF translocase^76,77^ - strictly require ssDNA accumulation beyond physiological levels expected at unperturbed forks. In this respect, an ATR-dependent mechanism to generate ssDNA via active resection of nascent DNA, while contributing to further ATR activation, may also represent a potent and tunable driver for the rapid extension of fork slowing and remodelling as a global, nuclear response (Fig. 6). In line with this concept, upon localized DNA damage, ATR-dependent phosphorylation of NBS1 on S343 - likely indicative of an activated MRN exonuclease, based on our data - is present not only at lesion sites but also throughout undamaged chromatin. Consistently, MRN exonuclease activity and the SMARCAL1 translocase are both needed for global replication fork slowdown.

Although nascent strand degradation (NSD) has been thoroughly studied downstream of fork reversal, it can also be observed upstream^32^. Of note, most studies investigating MRE11-driven NSD at forks have used comparably high levels of DNA damage or prolonged genome-wide fork stalling, where fork uncoupling *per se* may provide sufficient ssDNA to drive fork slowing and remodelling, bypassing the role of nucleolytic processing upstream of fork reversal^7^. A similar paradox has been described for RAD51, which acts both upstream and downstream of fork reversal, with the upstream function not requiring stable nucleoprotein filaments and being uncovered only upon full inactivation of the protein^11,78,12^.

IDRs and their post-translational modifications are rapidly emerging as key regulators of the DNA damage response^79^. Their known propensity to promote biomolecular condensates is proposed to complement more canonical lock-and-key interactions, providing a new layer of regulation for the multistep reactions mediating DNA damage signalling and repair ^46^. IDR-mediated regulation has been recently reported for multiple DNA damage response proteins, controlling the activity of MRN and other nuclease at both DSBs and stressed replication forks^46,80,81,82^ and offering new perspectives for therapeutic targeting^83^. Current structural and molecular information does not yet support a definitive mechanistic model for how NBS1 S343 phosphorylation may regulate MRN function in the context of active fork slowing. A vast protein portion at the C-terminus of NBS1 – including the IDR hosting S343 – is intrinsically disordered and flexible and was hypothesized to modulate intra-and intermolecular interactions that promote different functions of the MRN complex^84^. In particular, the IDR containing S343 is located in-between the N-terminal FHA-BRCT domains, which are important for phosphoprotein interactions (e.g. with CtIP and the MRE11-interacting domain at the C-terminus)^85,84^. We hypothesise that S343 phosphorylation may directly or indirectly mediate molecular interactions required for MRN activation in the context of replication factories, which may entail rather different regulatory mechanisms from those well-characterized in the context of DSB repair. The finding that MRN is already tightly associated with unperturbed replication forks^27,28,30^ supports the hypothesis that a rapid and intrinsic MRN activation mechanism could bypass the need for the recruitment of cofactors and quickly propagate the RS response throughout the nucleus. Notably, NBS1-S343 phosphorylation had been previously implicated in promoting chromatin accumulation of ATR and RPA phosphorylation upon prolonged fork stalling by HU^86^, highlighting the relevance of this residue in orchestrating yet-elusive mechanisms of the RS response. Thorough biochemical and structural investigations will need to clarify at the molecular level how S343 phosphorylation on NBS1 promotes MRN exonuclease activity in response to low dose RS, particularly in the context of DNA substrates that mimic low replication interference or even unperturbed replication forks.

Using systemic, low-dose UV-C irradiation it has been shown that NBS1 phosphorylation on S343 is ATR-dependent^69^, while the same phosphorylation event is fully ATR-independent and instead triggered by ATM upon ionizing irradiation^69^. Hence, the responsible kinase and the functional relevance of NBS1-S343 phosphorylation are dictated by the physiological context. In line with this, our DNA fiber assays revealed that replication fork slowdown in response to mild genotoxic treatments does not require ATM activity but relies on full ATR activity. Although we identified NBS1 phosphorylation as a necessary non-canonical ATR modification controlling fork progression under genotoxic stress, our data do not exclude the contribution of additional ATR-dependent events to this process. Elucidating the function of additional targets identified in our phospho-proteomic screen will be essential to refine this model in the future.

The importance of the MRN proteins in stress signalling mechanisms - even beyond replication forks and DNA repair - is highlighted by its recently described unexpected role as a direct stimulator of cGAS^87,88^. Our data extend the role of the MRN complex well beyond stalled forks and pathological conditions of nascent strand degradation^18,23,89^, identifying a crucial regulatory role for MRN-dependent DNA processing in modulating replication fork progression and architecture upon conditions that are compatible with residual fork progression. As mild replication interference is more likely to reflect clinically relevant conditions of anticancer treatments, a deeper investigation of these molecular events is likely to uncover actionable targets to potentiate cancer therapy. In this respect, it is encouraging that S343 phosphorylation is not essential for the key DNA damage response functions of the MRN complex^51^, while targeting this event showed promising result enhancing chemosensitivity of BCR/ABL-positive leukemia cells^90^. Further investigation of MRN regulation upon mild replication interference may thus represent an important avenue of research for future clinically relevant studies on the RS response.

## Material and Methods

### Cell lines

All Human osteosarcoma (U2OS) cells, Retinal pigmen epithelium (RPE-1) and Human colon cancer (HCT116) cells were cultured in DMEM (41966-029, Life Technologies) supplemented with 10 % fetal bovine serum (FBS, GIBCO), 100 U/ml penicillin and 100 mg/ml streptomycin in an atmosphere containing 5 % CO2 at 37°C.

U2OS Flip-In T-REx cells were grown in the DMEM medium supplemented with 10 % fetal calf serum (FCS, Gibco), 100 U/ml penicillin/streptomycin, 50 μg/ml hygromycin B and 10 μg/ml blasticidin S.

### U2OS Flip-In T-Rex – cell line generation

To introduce mutations of interest into plasmids site-directed mutagenesis was used. Primers for mutagenesis (GC-content 40–60%, GC-clamp, 18–30 bp, melting between 60–70°C, mismatched base/s toward the middle of primer) were designed using SnapGene. Mutations were introduced by PCR with the parameters indicated in the Phusion Polymerase protocol (NEB). For a 20 μl PCR reaction, 4 μl 5x HF buffer, 0.4 μl 10 mM DNTP, 2 μl 10 μM forward primer, 2 μl 10 μM reverse primer, 0.6 μl DMSO, and 0.2 μl Phusion polymerase were mixed with 10 ng DNA and filled up to 20 μl with H2O. After the PCR was completed 5 μl of the product was mixed with 3 μl Gel Loading Dye Purple (6x) (B7025S, New England Biolabs) and loaded onto an agarose gel (1% Agar diluted in 1x TBE) for gel electrophoresis. Gel electrophoresis was carried out at 100 V. The gel was then imaged using Quantum (Vilber Smart Imaging) and analysed for correct plasmid size. The residual PCR product was digested by the addition of 1 μl DPN1 restriction enzyme for 1 h at 37°C to be ready for bacteria transformation and construct amplification.

For bacteria transformation, 50 μl aliquots of Electrocompetent E. Coli stored (ThermoFisher) were used. The bacteria were first defrozen on ice for 15 min and 1 μl of previously mutated plasmid was added to each. The mixtures were left on ice for 20 min before being heat shocked for 45 sec at 42 °C. Afterwards, they were moved back on the ice for 2 min. A total of 700 μl of LB-media was added to each Eppendorf tube before they were left shaking for 1 h at 37°C. During shaking agarose, 60 mm Petri dishes with plasmid-matching antibiotic resistance were pre-warmed in an incubator at 37°C. 260 μl of the mixture was plated on each agarose plate and left overnight in an incubator at 37°C. The next day, single colonies were picked and transferred into falcon tubes with 5 ml LB-media and plasmid-matching antibiotic 1:1000. Falcons were left rotating at 37°C overnight. Then, constructs were expanded using a QIAprep Spin Miniprep Kit (27106, QIAGEN) according to manufacturer protocol. The concentration of the expanded plasmids was determined via Nanodrop (Thermo Scientific). The plasmids were then sent to Microsynth for sequencing with a matching custom primer (pResiduesNBS1seqFw: 5’-TTG GCG GTG ATT TTC ATG AC-3’) to validate the incorporation of mutations.

Plasmids with correct incorporation of intended mutations were either used directly or expanded using QIAGEN Plasmid Maxikit (12163, QIAGEN) according to manufacturer protocol. Cells were transfected at 70% confluence using Lipofectamine 3000 (L3000008, Thermo Fisher) by Flp-In™ T-REx™ Core Kit manufacturer instruction. After 24 h, the cells on each Petri dish were trypsinised and split into three wells of a 24-well plate. They were seeded at 70 % confluence per well in complete media (without antibiotics). Later after 24h 250 μg/ml of hygromycin B and 10 μg/ml of blasticidin S. was added. The medium was replaced every 3 days, and cells were selected for approximately 3 weeks. Resistant colonies were picked, expanded, and then characterized for inducible NBS1myc expression both by immunofluorescence microscopy and immunoblotting. To induce the expression of NBS1myc, cells were treated with 10nM doxycycline (DOX) for 24 h.

### RNAi experiments

For RNAi experiments, U2OS cells were transfected with the indicated siRNAs for 48 or 72 hours using RNAiMax (Thermo Fisher Scientific) according to manufacturer’s instruction.

siLuc (5’-GGAGGAAGAUGUCAAUGUUdTdT-3’; Microsynth)

siNBS1 (5’-GGAGGAAGAUGUCAAUGUUdTdT-3’; Microsynth)

siSMARCAL1 (5’-GCUUUGACCUUCUUAGCAAdTdT-3; Microsynth)

### Generation of Halo-SMARCAL1 knock-in cell line

All cloning procedures were performed using in-vivo assembly (IVA) cloning, as described^91^. Endogenous Halo–SMARCAL1 knock-in cell lines were generated using CRISPR–Cas9–mediated genome editing. A ∼1 kb homology arm spanning approximately 500 bp upstream and downstream of the SMARCAL1 start codon was amplified by PCR using KOD Hot Start DNA polymerase (Novagen, 71086-3) or Phusion High-Fidelity DNA polymerase (New England Biolabs, M0530S) and cloned into the pJET vector (Thermo Fisher Scientific, K1231). The resulting plasmid was linearized at the SMARCAL1 start codon by PCR, and a BSDR–P2A–HaloTag cassette amplified from an in-house plasmid was inserted by IVA cloning to generate the donor plasmid for N-terminal tagging. The insertion disrupts the sgRNA target site to prevent Cas9 re-cleavage after homology-directed repair. sgRNAs targeting the SMARCAL1 start codon were designed using CRISPR knock-in design tools (Horizon Discovery and IDT) and cloned into px330-U6-Chimeric_BB-CBh-hSpCas9 (Addgene #42230) following the protocol described^92^. CRISPR–Cas9 and donor plasmids were co-transfected into HCT116 OsTIR1(F74G) cells using FuGENE HD (Promega, E2311). Cells were selected with blasticidin, and resistant populations were single-cell sorted into 96-well plates and expanded. Correct integration of the Halo–SMARCAL1 cassette was verified by western blotting to confirm expression of the tagged protein and to assess expression levels relative to parental cells. Clones with correct integration and physiological expression levels were expanded for functional validation.

### Protein extraction and western blotting

Extracts from all cell lines were prepared in Laemmli sample buffer (4 % SDS, 20 % glycerol, and 120 mM Tris- HCl, pH 6.8). Equal amounts of protein (30–50 μg) were loaded onto 4 %–20 % Mini-PROTEAN TGX Precast Protein Gels (BioRad). Proteins were separated by electrophoresis at 160 mA followed by transferring the proteins to Immobilon-P membranes (Thermo Fisher Scientific) for 1 hour at 350 mA (4 °C) in transfer buffer (25 mM Tris and 192 mM glycine) containing 10 % methanol. Before addition of primary antibodies, membranes were blocked in 5 % milk in 0.1 % TBST (1 × TBS supplemented with 0.1 % Tween 20) for 1 hour and incubated in 3 % BSA with primary antibodies overnight at 4°C. Membranes were probed for the indicated antibodies. Secondary antibodies were added for 1 hour at room temperature (in blocking solution). Membranes were washed three times with 0.1 % TBST, 10 min each, after primary and secondary antibody incubations and detected with ECL detection reagent (GE healthcare).

### DNA fiber spreading assay

All cell lines subjected to this analysis were grown asynchronously, pre-treated as indicated in the corresponding Figures and labeled with 30 μM of the thymidine analog chlorodeoxyuridine (CldU; Sigma-Aldrich) for 30 min, they were then washed three times with warm PBS and subsequently exposed to 250 μM of 5-iodo-2ʹ-deoxyuridine (IdU) for 30 min alone or in combination with mild doses of genotoxic treatments (100 nM CPT, 20 nM ETP). All cells were collected by standard trypsinization and resuspended in cold PBS at 3.5 × 105 cells/mL. 3 μL of this cell suspension were then mixed with 7 μL of lysis buffer (200 mM Tris-HCl, pH 7.5, 50 mM EDTA, and 0.5% [w/vol] SDS) on a glass slide. After an incubation of 9 mins at RT, the slides were tilted at a 45° angle to stretch the DNA fibers onto the slide. The resulting DNA spreads were air-dried, fixed in 3:1 methanol/acetic acid, and stored at 4°C overnight. The DNA fibers were denatured by incubating them in 2.5 M HCl for 1 hour at RT, washed five times with PBS and blocked with 2 % BSA in PBST (PBS and Tween 20) for 40 minutes at RT. The newly replicated CldU and IdU tracks were stained for 2.5 hours at RT using two different anti-BrdU antibodies recognizing CldU (Abcam, ab6326, 1:500) and IdU (Becton Dickinson, 347580, 1:100), respectively. After washing five times with PBST (PBS and Tween 20) the slides were stained with Anti-mouse Alexa 488 (Invitrogen, A-11001, 1:300) and anti-rat Cy3 (Immuno Research, 712-166-1530, 1:150) secondary antibodies for 1 hour at RT in the dark. The slides were mounted in 30 μL Prolong Gold antifade reagent (Invitrogen). Microscopy was done using a Leica DM6 B microscope (HCX PL APO 63x objective). To assess fork progression the IdU/CldU ratio or IdU track lengths of at least 100 fibers per sample were measured using the line tool in ImageJ64 software. Statistical analysis was carried out using GraphPad Prism 7.

### Electron microscopic analysis of genomic DNA

Asynchronous and subconfluent cells (cell lines and genotypes are depicted in the corresponding figure panels and legends) were pre-treated as indicated in the respective figure panels. Cells were collected, resuspended in ice-cold PBS and crosslinked with 4,5ʹ,8-trimethylpsoralen (10 μg/ml final concentration), followed by irradiation pulses with UV 365 nm monochromatic light (UV Stratalinker 1800; Agilent Technologies). For DNA extraction (Muzi-Falconi and Brown, 2018), cells were lysed (1.28 M sucrose, 40 mM Tris-HCl [pH 7.5], 20 mM MgCl2, and 4% Triton X-100; Qiagen) and digested (800 mM guanidine–HCl, 30 mM Tris-HCl [pH 8.0], 30 mM EDTA [pH 8.0], 5% Tween-20, and 0.5% Triton X-100) at 50 °C for 2 h in presence of 1 mg/ml proteinase K. The DNA was purified using chloroform/isoamylalcohol (24:1) and precipitated in one volume of isopropanol. Finally, the DNA was washed with 70 % EtOH and resuspended in 200 μl TE (Tris-EDTA) buffer. 120 U of restriction enzyme (PvuII high fidelity, New England Biolabs) were used to digest 6 μg of the purified genomic DNA for 5 h at 37C. RNase A (Sigma– Aldrich, R5503) to a final concentration of 250 ug/ml was added for the last 2 h of this incubation. The digested DNA purified using the Thermo Fisher Silica Bead Gel Extraction kit according to manufacturer’s instructions. The Benzyldimethylalkylammonium chloride (BAC) method was used to spread the DNA on the water surface and then load it on carbon-coated 400-mesh nickel grids (G2400N, Plano Gmbh). Subsequently, DNA was coated with platinum using a High Vacuum Evaporator BAF060 (Leica). The grids were imaged automatically using a Talos 120 transmission electron microscope (FEI; LaB6 filament, high tension ≤120 kV) with a bottom-mounted CMOS camera BM-Ceta (4000x4000pixel) and the MAPS software package (Thermo Fisher Scientific, Eindhoven, The Netherlands) as described. Samples were annotated for replication intermediates using the MAPS Viewer software, overlapping images for annotated regions were stitched together using the automated pipeline ForkStitcher and final images were analysed using ImageJ. For each experimental condition at least 70 replication fork molecules were analysed in two different biological replicates.

### FACS analysis

For the analysis by FACS, cells were pulse-labeled with 10 μM 5-Ethynyl-2’-deoxyuridine (EdU, A10044, Thermo Fisher) for 30 min before trypsinization. After washes with 1x PBS, cells were incubated in 300 μl 4 % paraformaldehyde diluted in 1x PBS in 1.5 ml Eppendorf tubes for 15 min at room temperature. Afterwards, 700 μl 1% BSA diluted in 1x PBS was then added and the mixture was spun down at 1500 rpm in a microcentrifuge. After the supernatant was removed cells were washed again using 1 ml 1% BSA. Cells were spun down at 1500 rpm and then permeabilized in 100 μl 1x PBS with 10 % saponin buffer for 30 min at room temperature. After spinning down, they were then incubated with primary antibodies in 10 % saponin buffer in the dark for 1.5 h at room temperature. The primary antibody was removed by centrifugation, and cells were washed with 400 μl 10% saponin buffer and spun down at 1500 rpm. 50 μl 10 % saponin containing the secondary antibodies was then added and cells were incubated for 30 min in the dark at room temperature before being centrifuged. Cells were washed again using 400 μl 10 % saponin buffer and spun down. EdU was made detectable by 150 μl laboratory-made click-it reaction (100 mM Tris pH 8.0, 100 mM CuSO4, 20 mg/ml sodium-l-ascorbate, 10 mM azide). Cells were then washed by adding 400 μl 0.5 % BSA (diluted in PBS) and spun down. They were then resuspended in 0.5 % BSA containing RNase A (0.1 mg/ml) and DAPI (1 ug/ml) for 30 min at room temperature in the dark. The cells were then transferred to FACS tubes and examined using the Attune NxT Flow Cytometer (Invitrogen). Analysis and visualization of data were done using the FlowJo software (BD Biosciences).

### Phospho-proteomic screen

#### Fluorescence activated cell sorting

Approximately 10 M U2OS Fucci cells were harvested at 70 % confluence, and the pellet was resuspended in 1 ml of PBS. All sorting experiments were performed using BD FACS Aria III flow Cytometer (BD Biosciences), using the gating strategy illustrated in Fig. 2b (S phase cells). Cells were sorted at 37 °C with a 100 uM nozzle at an event rate not exceeding 3000 evts/s into batches of 1 M cells at a time. Immediately post sorting, the sorted batch was split into two samples of 500 k cells each and put back in culture. One sample was treated with ATRi (AZ20, 1 uM) and the other one left untreated for 1 hour. Following the optional pre-treatment with ATRi, both batches were treated with 20 nM ETP for another hour. All cells were then washed with PBS, pelleted and snap-frozen. A total of 50 M cells (e.g. 50 batches of 1 M sorted cells) were used for each treatment condition. Two independent biological replicates have been performed.

#### Phospho-proteomic data generation

Cell pellets were lysed in modified RIPA buffer as previously described^61^. Lysates were subject to tryptic digestion as previously described^93^. Tryptic digests were acidified and desalted using 200mg SepPak C18 cartridges. Phosphopeptide enrichment was performed using High Select Fe-NTA phosphopeptide enrichment kits following manufacturer protocols. Phosphopeptides from each condition were labeled with three TMT channels each from one kit of TMT sixplex. Labeling was performed as previously described^93^. After TMT labeling, pooled samples were pre-fractionated into 30 fractions via offline HILIC with a Dionex Ultimate 3000, as previously described^93^. Each fraction was injected and analyzed on a Q-Exactive HF Orbitrap mass spectrometer as previously described^93^.

#### Phospho-proteomic data analysis

The Trans Proteomic Pipeline (TPP) version 6.0.0 was used for phosphopeptide identification and quantification. MS data were converted to mzXML using msConvert as packaged with the TPP, after which spectral data files were searched using the Comet search engine (v2021 rev 1). Peptide identifications were validated using PeptideProphet, phosphorylation site localization was performed using PTM Prophet, and TMT channel quantification was performed using Libra. Results from Libra were exported as tab-delimited files for further processing via R scripts as previously described^93^ . The phosphoproteomic data and results generated in this study were deposited to PRIDE (assigned accession number PXD075629; Token V0t2qnhXVck8).

### Biochemistry

#### Expression and purification of recombinant proteins

The human MRN (MRE11-RAD50-NBS1) complex was expressed with pTP17 (Tanya Paull, University of Texas at Austin), pFB-RAD50-FLAG and pTP36 (Tanya Paull, University of Texas at Austin), coding for his-tagged MRE11, FLAG-tagged RAD50 and untagged NBS1, respectively^94^. NBS1 S343A variant was prepared by mutating the wild type plasmid by QuikChange site-directed mutagenesis kit following manufacturer’s instructions (Agilent Technology) and using the primers listed here below. All MRN variants were expressed in Sf9 insect cells in SFX Insect serum-free medium (Hyclone) using the Bac-to-Bac expression system (Invitrogen), according to manufacturer’s recommendations. Frozen Sf9 pellets from 500 ml culture were resuspended in lysis buffer (50 mM Tris-HCl pH 7.5, 1:400 protease inhibitor cocktail [Sigma, P8340], 30 µg/ml leupeptin [Merck Millipore], 1 mM phenylmethylsulphonyl fluoride [PMSF], 0.5 mM β-mercaptoethanol, 0.1 % NP40) and incubated at 4°C for 20 min. Glycerol was added to a final concentration of 25 %, NaCl was added to a final concentration of 325 mM and the cell suspension was incubated at 4°C for 30 min. The cell suspension was centrifuged at 55,000 g at 4°C for 30 min. The soluble extract was incubated with ANTI-FLAG M2 resin (Sigma) at 4°C for 1 h. The resin was washed with FLAG wash buffer (50 mM Tris-HCl pH 7.5, 0.5 mM β-mercaptoethanol, 300 mM NaCl, 10 % glycerol, 1 mM PMSF) supplemented with 0.1 % NP40. Before elution, the resin was washed twice with FLAG wash buffer without NP40. Proteins were eluted using FLAG elution buffer (50 mM Tris-HCl pH 7.5, 0.5 mM β-mercaptoethanol, 300 mM NaCl, 10 % glycerol, 1 mM PMSF, 200 ng/μl FLAG peptide [GLPBIO]). Fractions containing high protein concentration as estimated by the Bradford assay were pooled, aliquoted, snap-frozen in liquid nitrogen and stored at -80°C. The MRN(S343A) mutant was purified in the same way.

pCtIP was expressed using the pFB-MBP-CtIP-his plasmid ^94^ and purified by affinity chromatography exploiting the N-terminal MBP-tag and the C-terminal his-tag. The MBP-tag was removed using PreScission Protease after the amylose purification step. To preserve the phosphorylation state of CtIP (pCtIP), Sf9 cells were treated with 50 nM Okadaic acid (APExBIO) for 3 h before cell harvesting. 1 μM camptothecin (Sigma) was also added to further activate protein phosphorylation cascade 1 h before cell harvesting ^94^.

**Oligonucleotides for mutagenesis use in this study**.

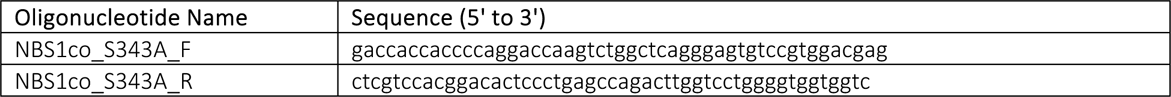

#### Preparation of oligonucleotide-based DNA substrates

All oligonucleotides were purified by polyacrylamide gel electrophoresis and purchased from Eurogentec. Oligonucleotides were radiolabeled at the 3’-end using Terminal Transferase (New England Biolabs) and [a-32P] dCTP (Hartmann Analytic) according to manufacturer’s instructions. Upon labeling, oligonucleotides were purified on a Micro Bio-Spin P-30 Gel Column (Bio-Rad). The labeled oligonucleotide was then annealed with a 2-fold excess of the respective unlabeled oligonucleotide in annealing buffer (10 mM Tris-HCl pH 8, 50 mM NaCl, 10 mM magnesium chloride), heated to 95°C for 3 min and cooled down to room temperature overnight. To prepare the 70-bp-long dsDNA substrate, PC210 oligonucleotide was labeled and annealed with PC211 oligonucleotide. The sequence of all oligonucleotides used in this study is listed below.

**Oligonucleotides use in this study**. The bold T represents the site of the biotin modification.

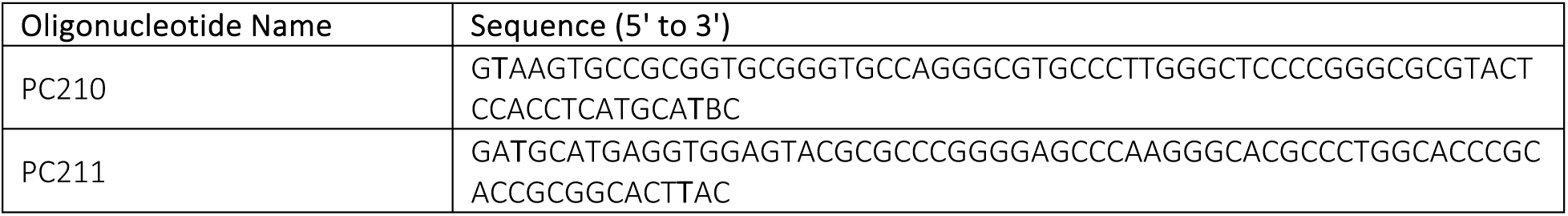

#### Electrophoretic mobility shift assay

The reactions (15 µl volume) were performed in binding buffer containing 25 mM Tris-acetate pH 7.5, 5 mM magnesium chloride, 1 mM manganese chloride, 1 mM DTT, 1 mM ATP, 0.25 mg/ml BSA (New England Biolabs) and 1 nM oligonucleotide-based DNA substrate (in molecules). After the addition of the MRN protein, the reactions were incubated on ice for 15 min. Loading dye (50% glycerol, bromophenol blue) was added and the products were separated by 4 % polyacrylamide (19:1 acrylamide-bisacrylamide, Bio-Rad) native gel electrophoresis in Tris-Acetate-EDTA (TAE) buffer. The gels were dried on a 17 Chr paper (Whatman) and exposed to storage phosphor screen (GE Healthcare) and scanned by a Typhoon Phosphor Imager (FLA 9500, GE Healthcare) and quantitated using ImageJ Software.

#### DNA endonuclease assays

Endonuclease assays (15 μl volume) were performed in nuclease buffer containing 25 mM Tris-acetate pH 7.5, 5 mM magnesium chloride, 1 mM manganese chloride, 1 mM dithiothreitol (DTT), 1 mM ATP, 0.25 mg/ml BSA (New England Biolabs) and 1 nM oligonucleotide-based DNA substrate (in molecules). Reactions were supplemented with 15 nM monovalent streptavidin and incubated for 5 min at room temperature to block the biotinylated ends of the DNA substrates. The recombinant proteins were then added to the reactions on ice and samples were incubated at 37°C for 2 h. Reactions were stopped by adding 0.5 μl ethylenediaminetetraacetic (0.5 M EDTA) and 1 μl Proteinase K (14-22 mg/ml, Roche), and incubated at 50 °C for 30 min. Finally, 16.5 μl loading buffer (5 % formamide, 20 mM EDTA, bromophenol blue) was added to all samples and the products were separated on 15 % polyacrylamide denaturing urea gels (19:1 acrylamide-bisacrylamide, Bio-Rad). The gels were fixed in fixing solution (40 % methanol, 10 % acetic acid, 5 % glycerol) for 30 min at room temperature and dried on a 3MM paper (Whatman). The dried gels were exposed to storage phosphor screen (GE Healthcare), scanned by a Typhoon Phosphor Imager (FLA 9500, GE Healthcare) and quantitated using ImageJ Software.

#### DNA exonuclease assays

Exonuclease assays with dsDNA substrate (15 μl volume) were performed in nuclease buffer containing 25 mM Tris-acetate pH 7.5, 5 mM manganese chloride, 1 mM DTT, 1 mM phosphoenolpyruvate (PEP), 80 U/ml pyruvate kinase, 0.25 mg/ml BSA (New England Biolabs) and 5 nM oligonucleotide-based dsDNA substrate (in molecules). The recombinant proteins were then added to the reactions on ice and samples were incubated at 37°C for 2 h. Reactions were stopped by adding 0.5 μl of EDTA (0.5 M) and 1 μl of Proteinase K (14-22 mg/ml, Roche), and incubated at 50 °C for 30 min. Finally, 16.5 μl of loading buffer (5 % formamide, 20 mM EDTA, bromophenol blue) was added to all samples and the products were separated on 15 % polyacrylamide denaturing urea gels (19:1 acrylamide-bisacrylamide, Bio-Rad). The gels were fixed in fixing solution (40 % methanol, 10 % acetic acid, 5 % glycerol) for 30 min at room temperature and dried on a 3MM paper (Whatman). The dried gels were exposed to storage phosphor screen (GE Healthcare), scanned by a Typhoon Phosphor Imager (FLA 9500, GE Healthcare) and quantitated using ImageJ Software.

#### Local UV-C irradiation through micropore filter

For local UV-C irradiation through micropore filters, cells were seeded onto glass coverslips (12 mm diameter, thickness No.1.5, Thorlabs; priorly coated with sterile 0.01 % Poly-L-lysine solution (Sigma-Aldrich)) and grown until 70 - 80 % confluency. On the day of the experiment cells were optionally pretreated with ATR or resection inhibitors (AZ20, 1 µM; 50 µM PFM39) for the indicated times and concentrations. Cells and micropore filters (Isopore Filter 5 um PC Membrane; 13 mm; TMTP01300) were then rinsed with warm PBS 1x and excess PBS was carefully removed from coverslips and filters right before placing the filters onto each coverslip. The filter-covered coverslips where then placed into a box with high walls to allow UV-C light solely coming from top through the pores of the filters. Exposure to UV-C light was carried out by a lamp (254 nm) for 3 seconds, corresponding to a dose of approximately 100 J/m^2^. Right after the UV-C pulse, fresh pre-warmed media was added to the wells, the filters removed and cells were allowed to recover for the indicated times, followed by an 8 mins pulse of 25 uM EdU incorporation. After the EdU pulse, cells were washed with cold PBS and pre-extracted with CSK buffer (10 mM Pipes pH7, 0.1 M NaCl, 0.3 M sucrose, 3 mM MgCl_2_, 0.5 % Triton-X) at 4°C for 4 mins. Cells were then carefully rinsed twice with cold PBS and fixed with 4 % PFA/PBS for 12 mins at RT. Fixation was followed by two rinses with PBS prior to denaturation (needed for the CPD antibody) with 0.07 M NaOH/PBS for 8 mins at RT. After washing with PBS, cells were permeabilized with Triton-X 0.3 %/PBS for 10 mins at RT. After two washes with PBS, samples were blocked with 5 % BSA/ 10 % FCS/ 1xPBS for a minimum of 1 h at RT. Primary antibody incubation was conducted in blocking buffer (anti-CPD 1:500, anti-pNBS1 1:500) over night at 4°C. Samples were washed twice with PBS and incubated with secondary antibodies (anti-mouse IgG2ak 555 and anti-rabbit 488, both 1:250) in blocking buffer for a minimum of 1 hour at RT in the dark. After two washes with PBS, cells were incubated with the EdU ClickIT reaction mix (85.5 mM TrisHCL, pH 8.0; CuSO_4_, 10 mM Sodium L-ascorbate, 5 uM azide 647) for 1.5 h at RT in the dark. After 3 washes with PBS, cells were incubated with DAPI in PBS for 15 mins, rinsed with PBS, mounted with Prolong Gold antifade reagent (Invitrogen) and allowed to dry prior to imaging using a Leica DM6 B microscope (HCX PL APO 63x objective).

Fluorescence images were acquired as single-plane multi-channel TIFF files and analyzed using ImageJ2 (software version: 2.16.0/1.54p) with a custom macro. Four channels were processed per field: DAPI (nuclei), pNBS1 or other (GFP), CPD (Cy3), and EdU (Cy5). Images were converted to 8-bit and background-subtracted using a rolling-ball algorithm. Nuclei were segmented from the DAPI channel following Gaussian blur (σ = 3) and Otsu thresholding with manual adjustment where required. Binary masks were generated and individual nuclei were defined using particle analysis. DNA damage regions were identified from the CPD channel after Gaussian blur (σ = 3), contrast enhancement (0.35 % saturation), and MaxEntropy thresholding. A CPD-positive mask defined damaged regions. Undamaged nuclear regions were generated by subtracting the CPD mask from the nuclear mask. For each nucleus, pNBS1 or other, CPD, and EdU signals were quantified in three compartments: whole nucleus, CPD-positive (damaged) regions, and CPD-negative (undamaged) regions. Mean fluorescence intensity was calculated as integrated density divided by the number of non-zero pixels within each region to avoid background bias. All images were processed using identical parameters across experimental conditions. Segmentation accuracy was verified by visual inspection prior to analysis and by measured CPD intensity levels (elevated in the damage region but undamaged regions comparable to nontreated samples).

#### Local UV-C laser micro-irradiation

For UVC laser micro-irradiation, cells were grown up to 70 % confluency on two glass coverslips (12 mm diameter, thickness No.1.5, Thorlabs), coated with sterile 0.01 % Poly-L-lysine solution (Sigma-Aldrich) and placed in the same 35 mm diameter plate (Falcon). One coverslip was subject to laser micro-irradiation, and the other one was used as a negative control. Cells were not directly grown on quartz coverslips because EdU incorporation was impaired on quartz (half less EdU incorporation compared to cells grown on glass coverslips, data not shown). Nuclei were stained by adding Hoechst 33258 (10 μg/mL final, Sigma-Aldrich) to the culture medium 15 min before UVC irradiation, then cells were rinsed with fresh medium.

The coverslip to be irradiated was flipped on a quartz coverslip (25 mm diameter, thickness No.1, SPI supplies) and transferred to a Chamlide magnetic chamber (Gataca-systems) on a custom stage insert (Live Cell Instrument) in a 37 °C heating chamber with CO2 supply. The control coverslip was also flipped on a quartz coverslip in the 35 mm diameter plate and kept in the same heating chamber.

Cells were irradiated for 120 ms using a 2 mW pulsed diode-pumped solid-state laser emitting at 266 nm (RappOptoElectronics, Hamburg GmbH) directly connected to a Zeiss LSM900 confocal microscope adapted for UVC transmission with all-quartz optics. The laser was attenuated using a 1 % neutral density filter and focused through a 40x/0.6 Ultrafluar glycerol objective with quartz lenses. The laser was controlled by a UGA Firefly module with SysCon2 software (RappOpto). The laser impact has an average size of 1 μm in diameter and damages around 1-2 % of the total nuclear volume. The corresponding UVC dose, not directly measurable but estimated by comparing the intensity of the CPD damage generated by the laser and the UVC lamp, is 1200 J/m2. Total irradiation time was 15 min to damage a minimum of 150 nuclei per experiment.

The glass coverslips were then flipped back in the original 35 mm diameter plate and after a 5 min recovery, the culture medium was replaced by fresh medium containing 25 µM Ethynyl-deoxy-Uridine (EdU, Invitrogen) for 10 min. Cells were pre-extracted with 0.5 % Triton X-100 in Cytoskeletal buffer (CSK: 10 mM PIPES pH 7.0, 100 mM NaCl, 300 mM sucrose, 3 mM MgCl2) for 5 min at room temperature to remove soluble proteins before fixation with 2 % paraformaldehyde (Electron Microscopy Sciences) for 10 min. Cells were subsequently denatured with 0,5 M NaOH/water for 5min RT, wash 3x with PBS and blocked with PBS-Tween 0,02 % + BSA 5 % for 15min RT. Primary antibody incubation (CPD (Kamiya Biomedical, #MC-062) 1/1000) was conducted in PBS-Tween/BSA 5 % for 1 h at RT. After 3 rinses with PBS-Tween secondary antibodies (goat anti mouse Alexa 568 (Invitrogen) 1/1000) were added for 45min to 1h at RT. After 3 rinses with PBS-Tween and one rinse with PBS, samples were fixed with 2 % PAF for 5min, rinsed 3x with PBS and permeabilized with PBS-triton 0,5 % for 15min at RT. After 3 rinses with PBS, cells were incubated with a ClickIT reaction mix (256 ul Buffer click it (kit), 12 ul CuSO4 (kit), 30 ul additive 1X (kit), 2 ul azide 488; 150 ul per coverslip) for 1h at RT. Cells were washed 3x with PBS and mounted in Vectashield with DAPI. Fluorescence imaging was performed using a Zeiss LSM900 confocal microscope equipped with a 40x/1.3 oil immersion Plan-Apochromat objective. Images were captured using Zen blue software version 3.5 and analyzed with ImageJ v.2.16.0/1.52.p (U.S. National Institutes of Health, Bethesda, Maryland, USA, http://imagej.nih.gov/ij/) as follows: The subtract background function was applied to all images prior to quantification. For segmentation purposes, images were smoothened using the median filter function. Nuclei were segmented based on Hoechst staining using the Default threshold function, and UVC-damaged regions were segmented based on CPD immunostaining using the Triangle or Li threshold function without manual adjustment and kept identical in each experiment. Masks of nuclei, damage, and damage-free nuclei were created and used to generate images containing damage-free nuclei or damage only using the Image Calculator function applied to the unsmoothened images. Mean intensities were measured in each region of interest and each channel by dividing raw integrated densities by the number of pixel values different from zero. Intensity profiles along a 1 µm wide line crossing the entire nucleus and passing through the damage area were obtained using the Plot Profile function in each channel.

**Key resources table**

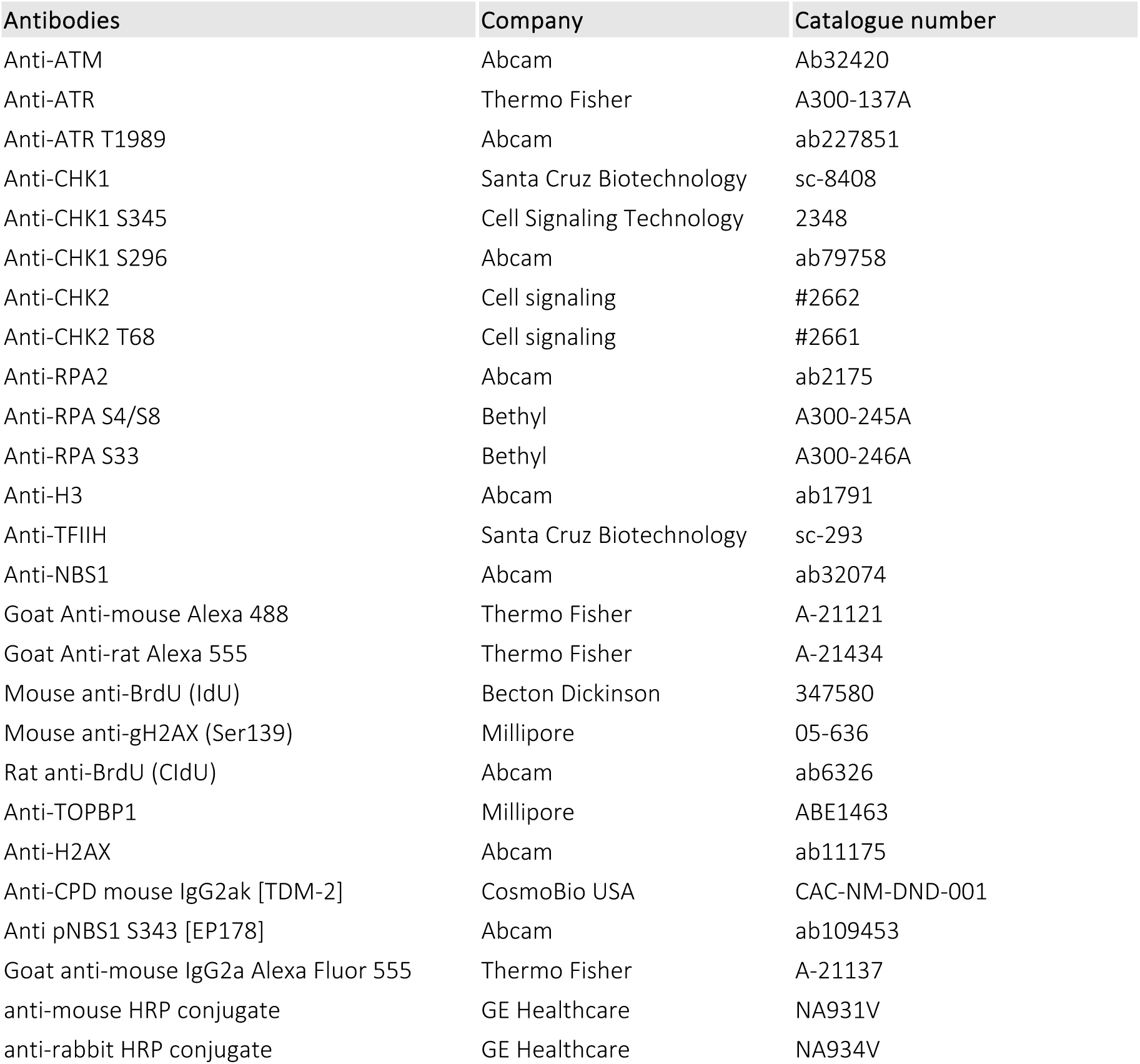
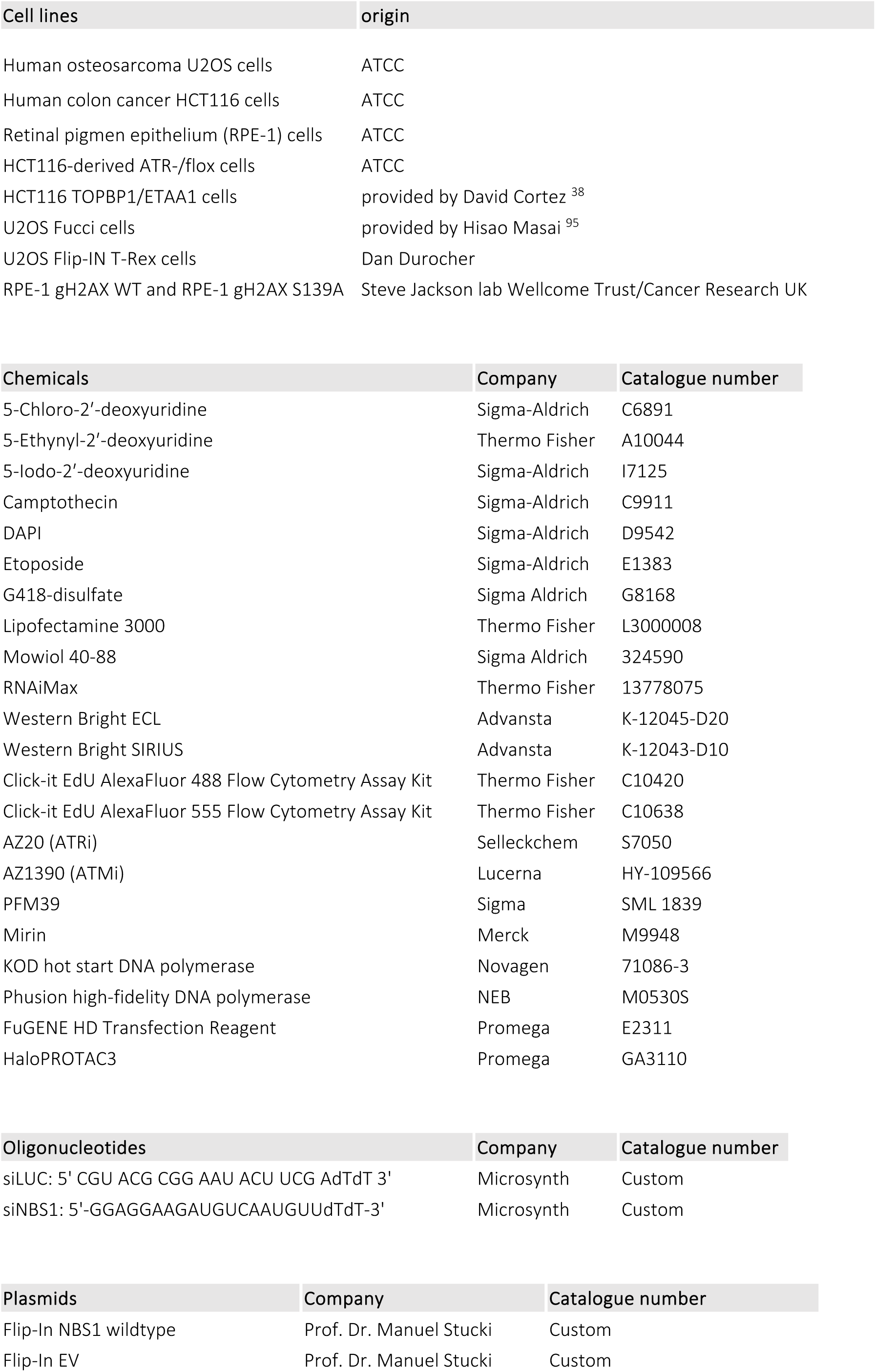
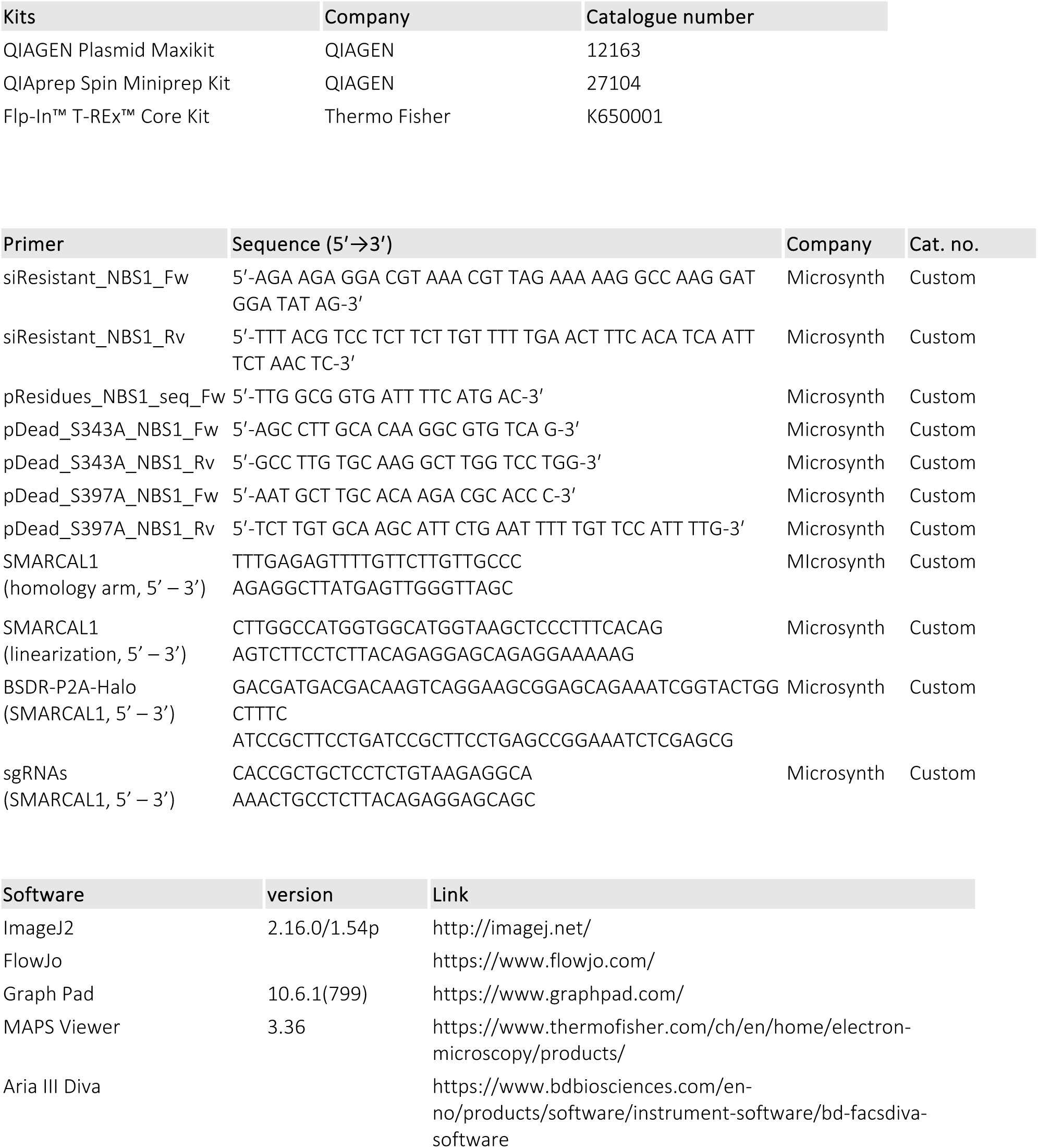

## Author contributions

M. L. conceived the project. J. K. and M. L. designed experiments, J. K. performed most experiments, data analysis, figure generation. M. L. and J. K. wrote the paper. W. J. C., D. D. and M. B. S. designed, conducted and analyzed the proteomics-screen. D. K. and M. A. performed EM experiments with MRE11 inhibitors. L. T. contributed in generating and validated the SMARCAL1-HP3 cells. I. C. and P. C., with the help of S. B., designed, conducted and analyzed all biochemistry experiments. S. Pi. and S. Po. designed and performed all laser-based micro-irradiation experiments. F. V. and A. A. S. designed and performed WB analysis of ATR/ ATM inactivation in Extended Data Fig. 1 c. All authors discussed the results, and reviewed the manuscript.

## Acknowledgements

We thank Lopes Lab members and Prof. Penengo for critical input on project design and realization, Dr. Daniel González Acosta for precious help with the Halo-SMARCAL1 cell generation. We thank David Cortez, Hisao Masai, Dan Durocher, Steve Jackson for making cell lines available, Manuel Stucki for providing us with NBS1 plasmids. We thank the UZH Center for Microscopy and Image Analysis and Cytometry Facility for their continuous support. Work in the Lopes lab has been supported by the Swiss National Science Foundation project grants 310030_189206 and 310030_219393, and by the Swiss Cancer League grant KLS-5698-08-2022. The Swiss National Science Foundation (SNSF) (Grants 310030_207588 and 320030-236167) and the European Research Council (ERC) (Grant 101018257) support the research in the Cejka laboratory. Work in the Smolka lab was supported by grants from the National Institute of Health, R35GM141159 to MBS, and F31CA281247 to WJC. Work in the Polo lab was supported by the European Research Council (ERC-2018-CoG-818625 “REMIND”) and an Emergence grant from Université Paris Cité (“DamSeg”). Work in the Sartori lab has been supported by the Swiss National Science Foundation project grant 310030_208143.

## Data availability statement

Raw data used to build all graphs and derive statistics - as well as original, uncropped blots – will be made available in a Source data file. Microscopy images are in the range of several Terabytes and would anyway require a trained eye for interpretation. They will hence be made available upon reasonable request.

## Competing interests

The authors claim no competing interests.

**Supplementary Table 1: List of non-canonical ATR targets identified in the phospho-proteomics screen (related to Figure 2 and Extended Data Fig. 2).**

Description of the individual columns as follows:

**Protein Phosphorylation Site:** Protein followed by residue numbers upon which phosphorylation sites are identified. “C” denotes a cluster of residues within which the phosphorylation site is located.

**Protein:** The protein upon which the phosphorylation event is identified.

**Peptide Sequence:** The amino acid sequence of the identified phosphopeptide with “phos” immediately after a phosphorylated residue or cluster or residues.

**Average Ratio:** The mean of the Log2 TMT ratios for each time the peptide was identified and quantified. Median Ratio: The median of the Log2 TMT ratios for each time the peptide was identified and quantified. Ratio SD: The standard deviation of the Log2 TMT ratios for each time the peptide was identified and quantified. Peptides: The number of times this peptide was identified and quantified.

**All ratios:** The Log2 TMT ratio for every time this peptide was identified and quantified.

**Best PTMProphet Localization Probability:** The best localization score calculated by PTMProphet from amongst the peptides identified and quantified.

**Flanking Sequence:** The 15 residues before and after the phosphorylated residue. For clusters and multiply phosphorylated peptides, this is shown for each phosphorylated residue.

---

**Extended Data Fig. 1:**
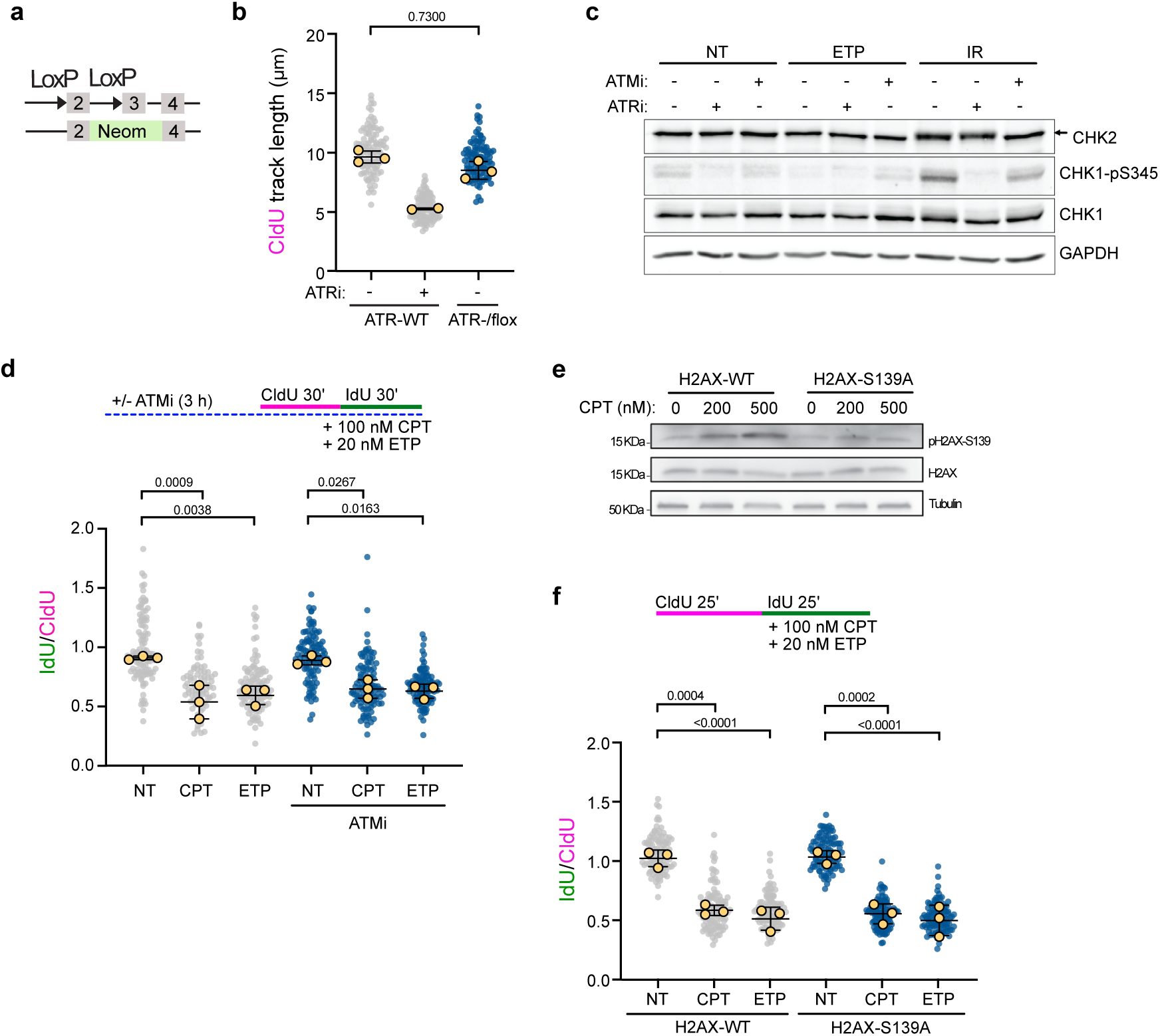
Full ATR activity is required for fork slowing and reversal, but canonical ATR activators, partners and targets are dispensable (related to Fig. 1). a, Schematic genotype of HCT116-derived ATR hypomorphic cells: One allele is disrupted by a Neomycine cassette, whereas on the intact allele, exon 2 is flanked bye loxP sites. b, Assessment of total CldU track length in CldU+IdU molecules (indirectly indicative of ground state replication fork speed) in ATR-WT versus ATR -/flox cells in experiments Fig. 1b-d. WT cells were optionally pre-treated with ATRi (AZ10, 1 uM, 1 h). c, Western blot analysis of nontreated, ETP-treated (20 nm, 1 h) and irradiated (IR, 5Gy, 1 h recovery) U2OS cells. Where indicated, cells have been pre-incubated with ATRi (AZ20, 1 µM, 1h pre-treatment, 2h total) or ATMi (AZ1390, 10 nM, 1h pre-treatment, 2h total). GAPDH serves as loading control. Black arrow indicates shift of the phosphorylated CHK2 protein. d, DNA fiber assay in U2OS cells, optionally pre-treated with an ATM kinase inhibitor (ATMi, AZ1390, 10nM, 1h pre-treatment, 2h total) and treated where indicated with 100nM CPT or 20nM ETP during the IdU pulse. top: Schematic treatment timeline. bottom: IdU/CIdU ratio is plotted for a minimum of 100 forks from a single representative experiment (indicated as grey/ blue dots). Overlayed yellow dots indicate medians from 3 individual biological replicates. Black line indicates the mean of the medians +/- SD. e, Western blot analysis of RPE-1 cells wild type (WT) for H2AX or mutated (S139A), treated where indicated with the depicted doses of CPT. Tubulin serves as a loading control. f, DNA fiber assay in RPE1 cells from d treated where indicated with 100nM CPT or 20nM ETP during the second label. top: Schematic treatment timeline. bottom: IdU/CIdU ratio is plotted for a minimum of 100 forks from a single representative experiment (indicated as grey/ blue dots). Overlayed yellow dots indicate medians from 3 individual biological replicates. Black line indicates the mean of the medians +/- SD. b, d, f, p-values were assessed using one-way ANOVA.

**Extended Data Fig. 2:**
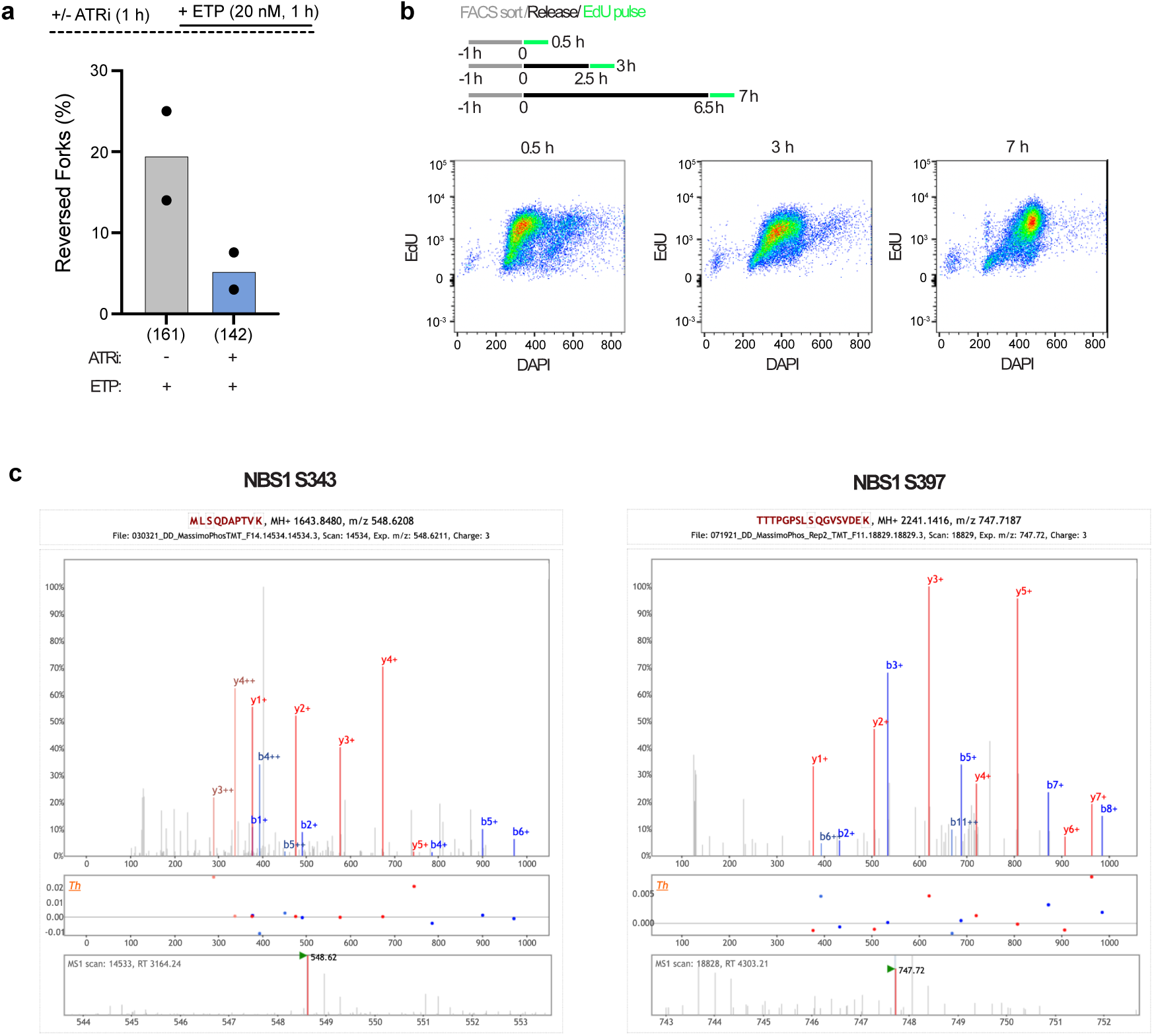
Phosphoproteomic screen validation (related to Figure 2). a, Frequency (in %) of reversed replication forks isolated from ETP (20nM, 1h) and optionally ATRi (AZ20, 1uM, 2h) treated U2OS cells. Bar graphs depict mean from two independent EM experiments (black dots, respectively). Total number of analyzed molecules in brackets. b, FACS analysis of actively replicating cells after sorting as described in Figure 2 b. Post-sort cells were allowed to recover in conditioned media for the indicated times, followed by an 0.5h EdU pulse and FACS staining. c, Annotated MS2 spectra for NBS1 phosphopeptides with decreased abundance following ATR inhibition; red: ETP spectra – blue ETP+ ATRi spectra.

**Extended Data Fig. 3:**
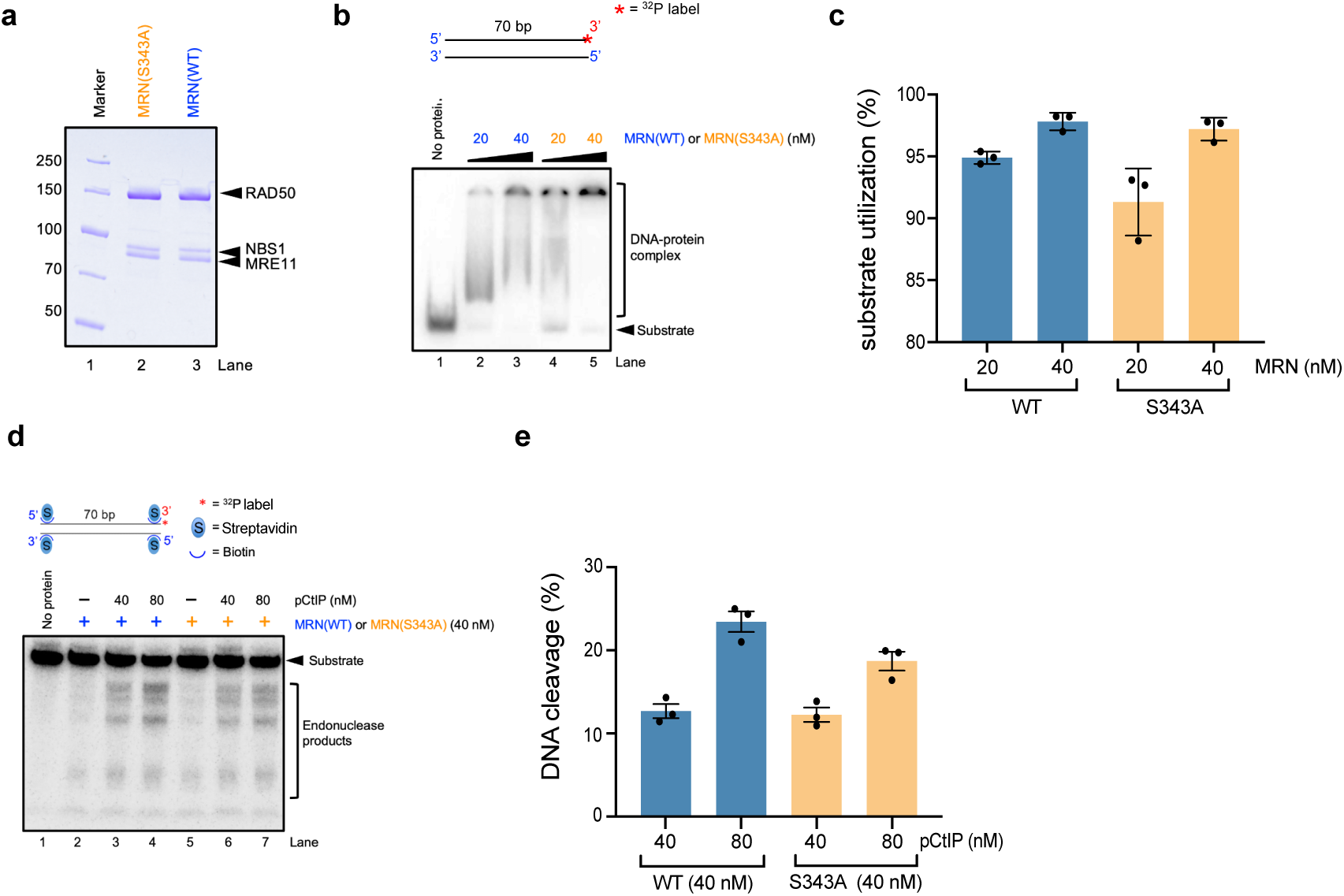
A single amino acid substitution in NBS1 (S343A) is permissive for MRN complex formation, DNA binding and endonuclease activity (related to Fig. 3). a, Recombinant MRN protein complex purified from Sf9 insect cells, composed of his-tagged MRE11, FLAG-tagged RAD50 and untagged NBS1, either as NBS1 wild type (in MRN – WT) or mutated for a single amino acid (in MRN-S343A). The polyacrylamide gels were stained with Coomassie Brilliant Blue. b, Electrophoretic mobility shift assay (EMSA) using a ^32^P-labeled double-stranded 70bp DNA substrate and the purified protein complex variants from a. top: Cartoon of the DNA substrate used for EMSA. bottom: Representative gel image visualizing the free and protein-bound dsDNA substrate. c, Quantification of substrate utilization from b. Bar graphs display the mean+/-SEM, black dots represent three independent replicates. d, Endonuclease assay using a ^32^P-labeled 70bp, double-stranded 70bp DNA substrate with biotinylated ends protected by streptavidin. top: Cartoon of the DNA substrate. bottom: Representative gel image visualizing endonucleolytic degradation by the indicated amounts of MRN protein complex variants, stimulated by recombinant phosphorylated CtIP (pCtIP). e, Quantification of endonuclease products from d. Bar graphs display mean+/- SEM, black dots represent three independent replicates.

**Extended Data Fig. 4:**
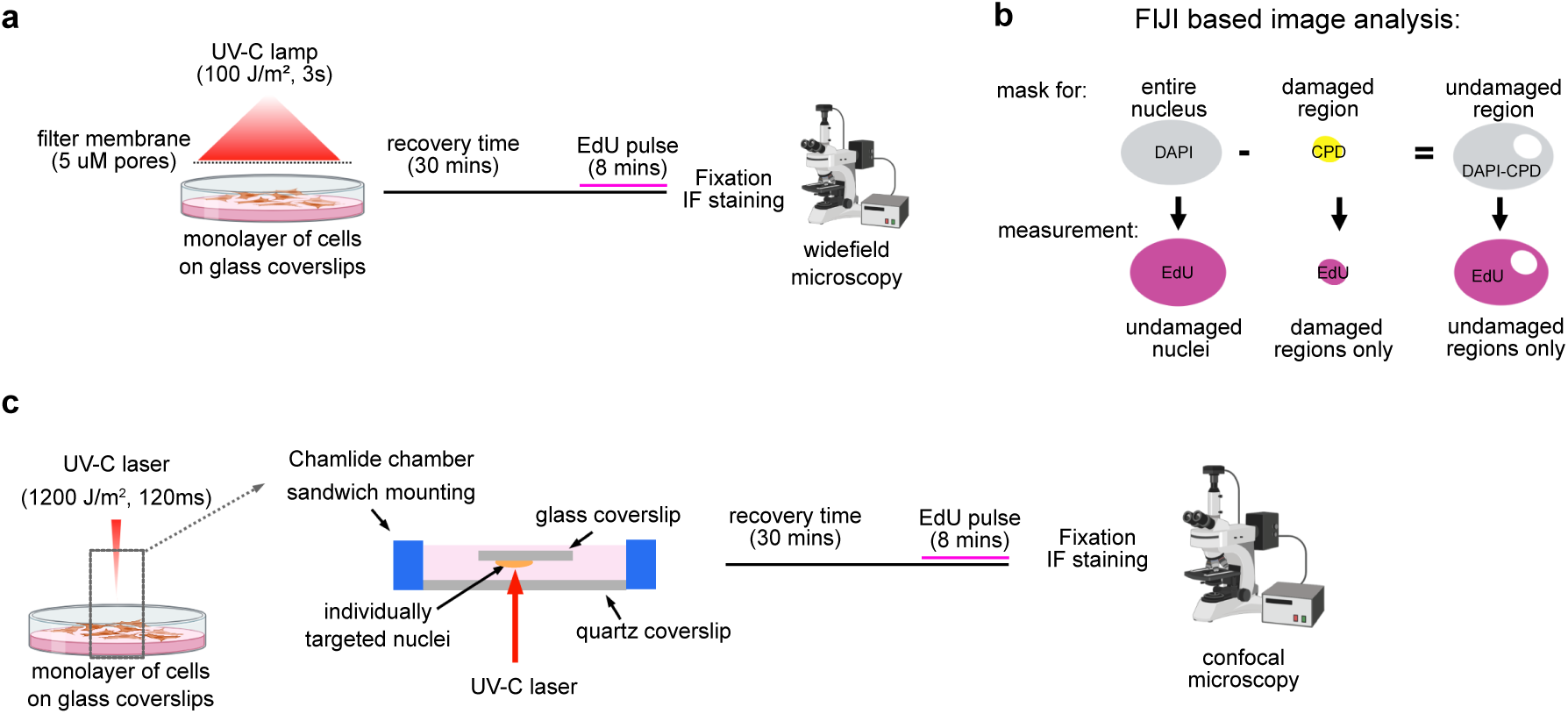
Experimental setups for localized UV-C irradiation (related to Fig. 4). a, Schematic setup used for micro-irradiation in Fig. 4a-e and 5a-f. UV-C irradiation (266nm, 100J/m^2^, 3s) was delivered to a subconfluent monolayer of cells grown on glass coverslips and shielded with a micropore filter (5µM pore size). Irradiated cells or a non-irradiated controls were allowed to recover for 30mins. A short EdU pulse was applied during the last 8mins of recovery, followed by pre-extraction, fixation, immunofluorescence staining and imaging with a widefield microscope. b, Schematic of simplified FIJI-based image analysis workflow used throughout Fig. 4-5. Entire nuclei were masked based on DAPI counterstaining, and damaged areas were defined using the CPD antibody signal. Substraction of the damaged region mask from the nuclear mask generated a mask for the undamaged region. All three regions were used to quantify the indicated mean intensities. c, Schematic setup of UV-C lacer micro-irradiation used in Fig. 4e-i: As in a, subconfluent cell monolayers grown on glass coverslips were used for UV-C irradiation. Herein, they were assembled in a Chamlide chamber sandwich mounting with the UV-C laser traversing the quartz coverslip at the bottom, reaching the targeted cells on an inverted glass coverslip. Post irradiation, the glass coverslip was turned upright again, left to recover for a total time of 30 mins, with a 10 mins EdU pulse at the end. Post pre-extraction, fixation and staining, cells were imaged with a confocal microscope.

**Extended Data Fig. 5:**
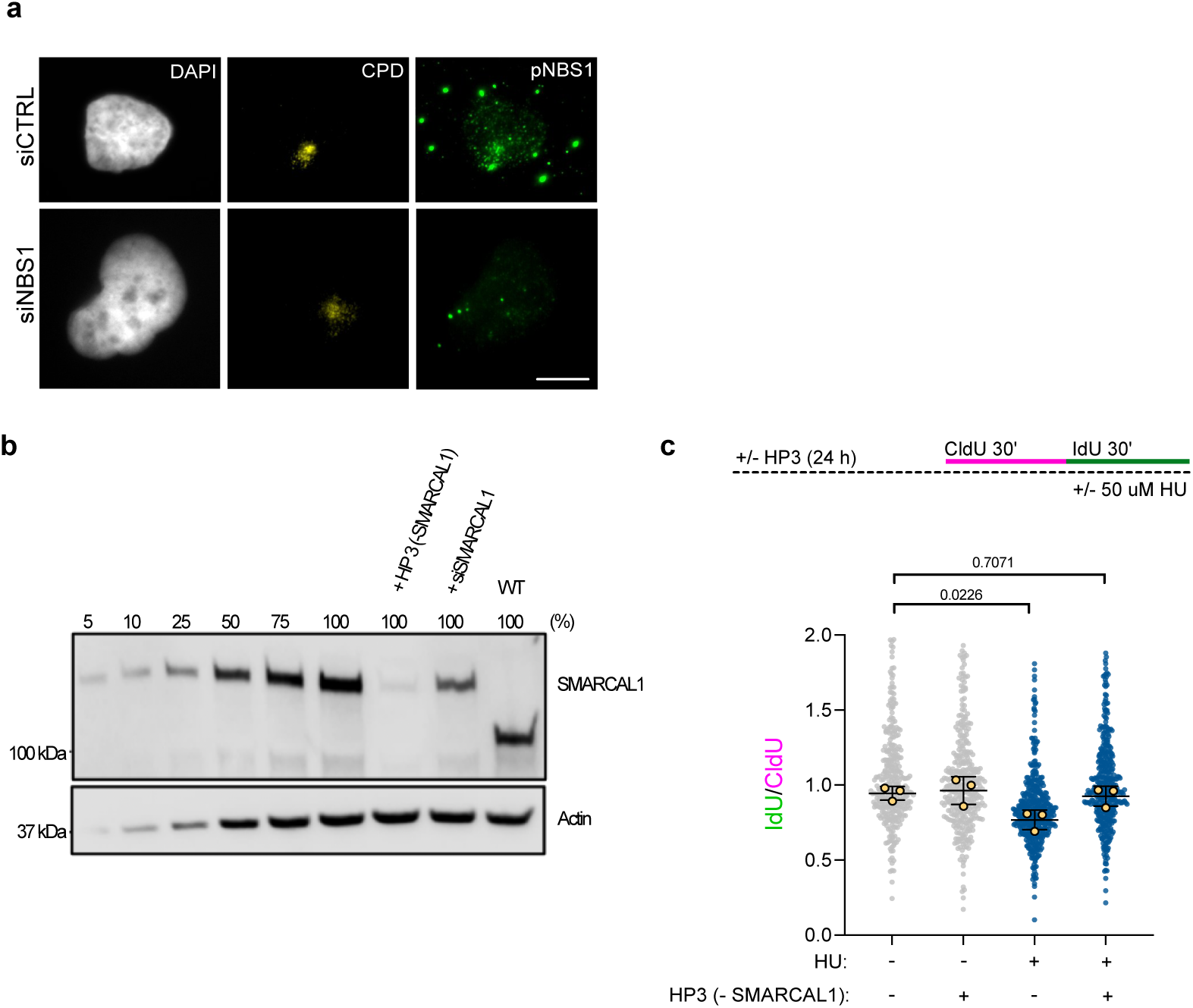
pNBS1 antibody and SMARCAL1 HaloPROTAC3 validation (related to Figure 5). a, To assess antibody specificity, U2OS cells were either transfected with siCTRL or siNBS1 72 h before UV-C irradiation as in Fig. 4a. and pNBS1 antibody staining as in Fig. 5c. b, Western blot analysis of SMARCAL1 protein levels and Actin as loading control in increasing protein amounts of a HCT116-derived SMARCAL1 HaloPROTAC3 cell line (in %). Cells were either left untreated, or, where indicated, SMARCAL1 was depleted using HaloPROTAC3 (+HP3) or an siRNA oligo directed against *SMARCAL1* (+siSMARCAL1). Parental HCT116 cells (WT) still contaning the endogenous (untagged) protein were loaded as comparison. c, DNA fiber assay in HCT116-derived SMARCAL1 HaloPROTAC3 cell from a, treated where indicated with 1µM HP3 for 24h and 50µM Hydroxyurea (HU) during the IdU pulse. top: Schematic treatment timeline. bottom: IdU/CIdU ratio is plotted for a minimum of 100 forks from a single representative experiment (indicated as grey/ blue). Overlayed yellow dots indicate medians from 3 individual biological replicates. Black line indicates the mean of the medians +/- SD. P-values were assessed with one-way ANOVA.

